# Genetic and environmental canalization are not associated among altitudinally varying populations of *Drosophila melanogaster*

**DOI:** 10.1101/715649

**Authors:** Maria Pesevski, Ian Dworkin

## Abstract

Organisms are exposed to environmental and mutational effects influencing both mean and variance of phenotypes. Potentially deleterious effects arising from this variation can be reduced by the evolution of buffering (canalizing) mechanisms, ultimately reducing phenotypic variability. As such, there has been interest regarding the plausible conditions that enable canalizing mechanisms to evolve. Under some models, the circumstances under which genetic canalization evolves is limited, despite apparent empirical evidence for it. It has been argued that canalizing mechanisms for mutational effects may evolve as a correlated response to environmental canalization (the congruence model). Yet, empirical evidence has not consistently supported the prediction of a correlation between genetic and environmental canalization. In a recent study, a population of *Drosophila melanogaster* adapted to high altitude showed evidence of genetic decanalization relative to those from low-altitudes. Using strains derived from these populations, we tested if they also varied for environmental canalization, rearing them at different temperatures. Using wing morphology, we quantified size, shape, cell (trichome) density and frequencies of mutational defects. We observed the expected differences in wing size and shape, cell density and mutational defects between the high- and low-altitude populations. However, we observed little evidence for a relationship between a number of measures of environmental canalization with population or with visible defect frequency. Our results do not support the predicted association between genetic and environmental canalization.

## Introduction

In addition to differences in trait means, there can be considerable variation in how much variation is observed among individuals of a given genotype (Waddington, 1942; Rendel, 1963; Félix and Barkoulas, 2015; Flatt, 2005; Visser et al., 2003; Siegal and Leu, 2014; Klingenberg, 2019; Mayer and Hansen, 2017; Pélabon et al., 2010; Gibson and Wagner, 2000). Theoretical work has examined this propensity to vary with respect to the evolution of phenotypic robustness or canalization. Such evolved properties are important to examine, as environmental and mutational variation influence trait variance, ultimately influencing organismal performance and fitness (Arnold, 2003, 1983). The sensitivity of a given genotype in its response to mutational or environmental influences can vary *among* genotypes. It has been empirically demonstrated that under mutational or environmental perturbation there is often the expression of cryptic genetic variation, which has previously been used as evidence for genetic canalization (Paaby et al., 2015; Paaby and Rockman, 2014; Scharloo, 1991; Gibson and van Helden, 1997; Dworkin, 2005a). Theory suggests that robustness to environmental variation — environmental canalization — can readily evolve as organisms are constantly exposed to the influence of environmental effects (Eshel and Matessi, 1998; Wagner et al., 1997). Yet, as deleterious mutations are often purged by natural selection, this can result in weak selection for genetic canalization (assuming stabilizing selection on the trait) making it potentially less likely to evolve (Wagner et al., 1997; Proulx and Phillips, 2005; Visser et al., 2003; Gibson and Wagner, 2000).

The congruence hypothesis was proposed as a solution to the apparent inconsistency between theoretical and empirical work regarding the evolution of genetic canalization (Wagner et al., 1997). This hypothesis predicts that genetic canalization evolves as a correlated response during selection for environmental canalization (Wagner et al., 1997; Gibson and Wagner, 2000; Visser et al., 2003). Empirical evidence for associations between genetic and environmental canalization is mixed. Some studies provide supporting evidence both from simulations (Ancel and Fontana, 2000; Siegal and Bergman, 2002; Shu et al., 2007) and empirical work (Lehner, 2010; Stearns and Kawecki, 1994; Stearns et al., 1995; Szöllősi and Derényi, 2009). However, some explicit tests for the congruence model did not find support for it (Dworkin, 2005a,c; Borenstein and Ruppin, 2006). The most likely explanation is that the evolution of genetic and environmental canalization are not homogeneous, given the complex interplay of selection, mutation rates, genetic architecture, and evolutionary history.

There are a number of important methodological and conceptual issues that influence the debate on the congruence hypothesis, and the study of the evolution of canalization more generally. First, the conditions in which the release of cryptic genetic variation can be used to infer genetic canalization may be more limited than once thought (Hermisson and Wagner, 2004; Geiler-Samerotte et al., 2019). Rather, mutation accumulation or mutagenesis experiments are likely to be more fruitful for investigating genetic canalization. Second, environmental canalization is often measured using multiple approaches that differ with respect to what aspects of environmental robustness they seek to capture. Within-individual variation (fluctuating asymmetry), among-individual within-genotype variation, and reaction norm of trait means under common or different environmental treatments (Dworkin, 2005b) have all been employed. Using these methods to study genetic and environmental canalization, some studies have seen modest evidence of association between degree of sensitivity to genetic perturbation (changes in trait means) and within- or among-individual variance within a genotype (Dworkin, 2005a,c; Chari and Dworkin, 2013; Camara and Pigliucci, 1999; Chandler et al., 2017). However, a number of other studies do not show a consistent relationship between magnitude of perturbation and among-individual within-line or within-individual variance (Haber and Dworkin, 2017; Debat et al., 2009; Levy and Siegal, 2008). This suggests there are multiple, partially distinct properties when considering robustness for a given genotype.

Surprisingly, one issue that has not been broadly considered in the literature regarding the evolution of canalization is the influences of both lab adaptation (domestication) and the use of lab induced mutations. Most studies of canalization and robustness use lineages likely to have undergone some degree of adaption to lab environments. Furthermore, many studies often use lab induced mutations as a source for genetic perturbations (Paaby et al., 2015; Gibson and van Helden, 1997; Dworkin, 2005a,c; Haber and Dworkin, 2017; Hallgrímsson et al., 2006; Debat et al., 2011; Levy and Siegal, 2008). However, lab domestication and induced mutations may be unrepresentative of natural populations (Dittmar et al., 2016; Rockman, 2008; Orgogozo et al., 2015). Lab induced mutations likely do not reflect the spectrum of mutational effects experienced by natural populations. This may bias inferences regarding the ability of genotypes to buffer the effects of mutations organisms are exposed to during their evolutionary history. Furthermore, when considering the evolution of canalization, the evolutionary history of the experimental populations matters. In some experimental studies, collections of natural lineages or families are used that have heterogeneous geographical origins, and/or have been maintained in the lab for long periods of time (i.e. Dworkin (2005a,c)). Thus, explicit tests for the correlated evolution of genetic and environmental canalization (*sensu* Wagner et al. (1997)) is difficult without knowledge of the evolutionary history of such populations.

Another shortcoming of many empirical studies examining properties of phenotypic robustness and canalization is that they have examined variation with a univariate perspective (Dworkin, 2005a,c), even when examining many traits (Levy and Siegal, 2008; Takahashi et al., 2011, 2010; Takahashi, 2017). Considerable evidence and theory have demonstrated that a multivariate perspective on evolutionary change 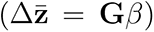 improves predictions and understanding of evolutionary responses to selection (Lande and Arnold, 1983; Lande, 1979; Schluter, 1996; Walsh and Blows, 2009; Houle et al., 2017; Blows and Mcguigan, 2014; Mcguigan and Blows, 2007; Pitchers et al., 2014; Agrawal and Stinchcombe, 2009; Hansen and Houle, 2008). Yet, this perspective has only been considered in a modest number of studies examining variational properties of phenotypes (Debat et al., 2009, 2011; Hallgrímsson et al., 2009, 2006; Green et al., 2017; Debat et al., 2006; Breuker et al., 2006; Cheverud et al., 1983; Pavlicev et al., 2009). When considering properties of trait (co)variation in this perspective, it is not just the magnitude of variation (matrix size), but direction (of major axes of variation) and the shape of the variance-covariance matrix (a proxy for trait integration) need to be considered as well. In a recent study examining variation in **E** (from **P** = **G** +**E**) across naturally derived strains and lab induced mutations, it was demonstrated that changes to trait means and relative orientation (directions of major axes of variation) of phenotypic (co)variances matrices were more variable than phenotypic integration (Haber and Dworkin, 2017). This suggests that a multivariate perspective needs to be consistently applied to studies examining trait (co-)variation (Klingenberg, 2019).

Thus what has been lacking for empirical studies testing evolutionary models of canalization is a system with the relevant natural history that can be studied with a multivariate approach. Lack et al. (2016a), among other recent studies, demonstrated that populations of *Drosophila melanogaster* from sub-Saharan Africa have recently adapted to a high-altitude environment. As is common for small insects evolving to high altitude environments (Dillon et al., 2006), they show increases in cold-tolerance (Pool et al., 2016), melanism (Bastide et al., 2016). They have also adapted via increased body size, wing size and shape (Pitchers et al., 2013; Lack et al., 2016a,b; Bastide et al., 2016; Fabian et al., 2015; Klepsatel et al., 2014), likely to deal with changes in flight response in cold, thin air (reviewed in Dillon et al. 2006). Intriguingly, there is a substantial increase in the frequency of qualitative mutational defects of wing morphology in the high altitude population (Lack et al., 2016a). Partially inbred Strains derived from a high-altitude Ethiopian population have defect frequencies as high as 40-50% (Lack et al., 2016a). This increase in frequency was not simply a result of hitchhiking of deleterious alleles, or a strong bottleneck, (Pool et al., 2012) but due to reduced mutational robustness, as assessed using mutagenesis experiments (Lack et al., 2016a). Importantly, the population-specific mutational sensitivities are pleiotropically linked to variants that influence the increase in wing size. This appears to be the case whether considering the variants among the high- and low-altitude populations (Lack et al., 2016a), or even in putative ancestral lowland populations (Groth et al., 2018) that has been artificially selected for large size. Currently, it is inferred that the increase in mutational sensitivity in the high-altitude population may have been a result of strong directional selection leading to rapid adaptation, with negative pleiotropic consequences. Compared with the ancestral low-altitude population that likely experienced a long history of stabilizing selection (and thus potentially promoting the evolution of canalization), the increase in size due to adaptation to conditions at high altitude, or due to strong artificial selection (Groth et al., 2018) have resulted in the evolutionary loss of canalization. This may represent a situation similar to that envisioned by Waddington (1942), where the population has not yet re-evolved its canalization mechanism after a long bout of strong directional selection for larger body size, wing size and shape.

The evolutionary history of the high- and low-altitude populations provides an ideal opportunity to test the relationship between genetic and environmental canalization. While there is considerable genetic variation within populations, strains derived from a high-altitude population in Ethiopia from an elevation of *∼*3000 m are much larger in body and wing size, have distinct wing shapes and have a greater frequency of qualitative ‘mutant’ phenotypes than low-altitude populations from Zambia from an elevation of *∼*500m (Lack et al., 2016a). Wing size and shape in *D. melanogaster* is a model system for studies of plasticity, sensitivity to mutational perturbation and within- and among-individual variability using both natural and lab-induced variation (Haber and Dworkin, 2017; Debat et al., 2011; Breuker et al., 2006; Debat et al., 2006; Soto et al., 2008; Klingenberg and Zaklan, 2000; Pélabon et al., 2006). In this study, we compared environmental canalization between the high and low-altitude populations. We used different measures of environmental canalization: within-line, among-individual variation (micro-environmental variation), within-individual variation (fluctuating asymmetry) and phenotypic plasticity across a temperature gradient (macro-environmental variation). Further, we correlated the different measures of environmental canalization and mutational perturbation in order to test the congruence hypothesis and determine whether strains with greater proportion of defects are also more variable (aka more decanalized). Despite demonstrating substantial population and environmental differences in wing size, shape, cell density and penetrance of mutational perturbation consistent with previous studies, we observed no consistent differences in measures of micro-environmental and macro-environmental canalization among populations. These results are discussed within the context of our ongoing understanding of the evolutionary mechanisms that influence trait variability.

## Materials and Methods

### Fly strains and Growth conditions

*Drosophila melanogaster* strains used in the current study represent a subset of those from Lack et al. (2016a,b). The high-altitude inbred strains were derived from flies collected in Fiche, Ethiopia at an altitude of 3070 m in December 2011. The low-altitude strains were collected in Siavonga, Zambia at an altitude of 530 m, and a 3125 km linear distance away from the high-altitude Ethiopian population in July 2010. These strains underwent inbreeding in the lab, which is expected to substantially reduce the effects of lab adaptation (because of the small *N_e_* within each strain), but this does result in substantial genetic drift within lines (but should not substantially alter allele frequencies among lines).

The flies for the micro-environmental variation experiments were raised as per Lack et al. (2016a). Flies were raised at 25°C, in 70% humidity, with 12:12 hour light/dark chamber, on standard cornmeal molasses food at a low larval density. This experiment was performed in June 2013.

A subset of the strains described above were used for the temperature plasticity (macro-environmental variation) and fluctuating asymmetry experiments. These strains were raised on a 1:1.5 protein to sugar ratio diet; recipe outlined in Table S3 at 24°C for two generations prior to the experiment. Newly emerged adults, 10-20 males and females, were collected and placed in egg collection chambers with apple agar plates with yeast. Eggs were collected, 50 at a time, and placed into vials with food. The flies were raised at 18°C, 24°C, and 28°C in 12:12 hour light/dark chambers until emergence. Adults were collected within 2 days of emergence and preserved in 70% ethanol. This experiment was performed in 2017. While these strains are partially inbred, as a check, we confirmed that the phenotypic effects of the size related traits from these lineages remained correlated with the low altitude populations (where variation for size is considerable), and also showed the same overall patterns (for mean size, shape and defect frequencies).

### Phenotyping

#### Wing size and shape - Micro-environmental canalization

The right wing of each fly was dissected and imaged using an Olympus DP30B camera mounted on an Olympus BX51 microscope (Olympus software V.3,1,1208) using a 2x objective (20X total magnification). Landmark and semi-landmark data was captured using a modified version of the “WINGMACHINE” pipeline (Houle et al., 2003; Pitchers et al., 2019). Coordinates of two starting landmarks were recorded using tpsDig2 software (V2.16). These coordinates are the humeral break on the leading edge of the wing and the alula notch on the trailing edge of the wing. B-splines were fit to veins and wing margin for each image using Wings (V3.7), reviewed and manually adjusted, if necessary. Landmark and semi-landmark positions were extracted and the shape information after adjusting for size, position, and rotation information using CPReader software (V1.12r). This produced data composed of 12 landmarks and 36 semi-landmarks (Figure 1B) as well as centroid size of for each specimen. The strains used in this experiment are outlined in Table S1.

**Figure 1:**
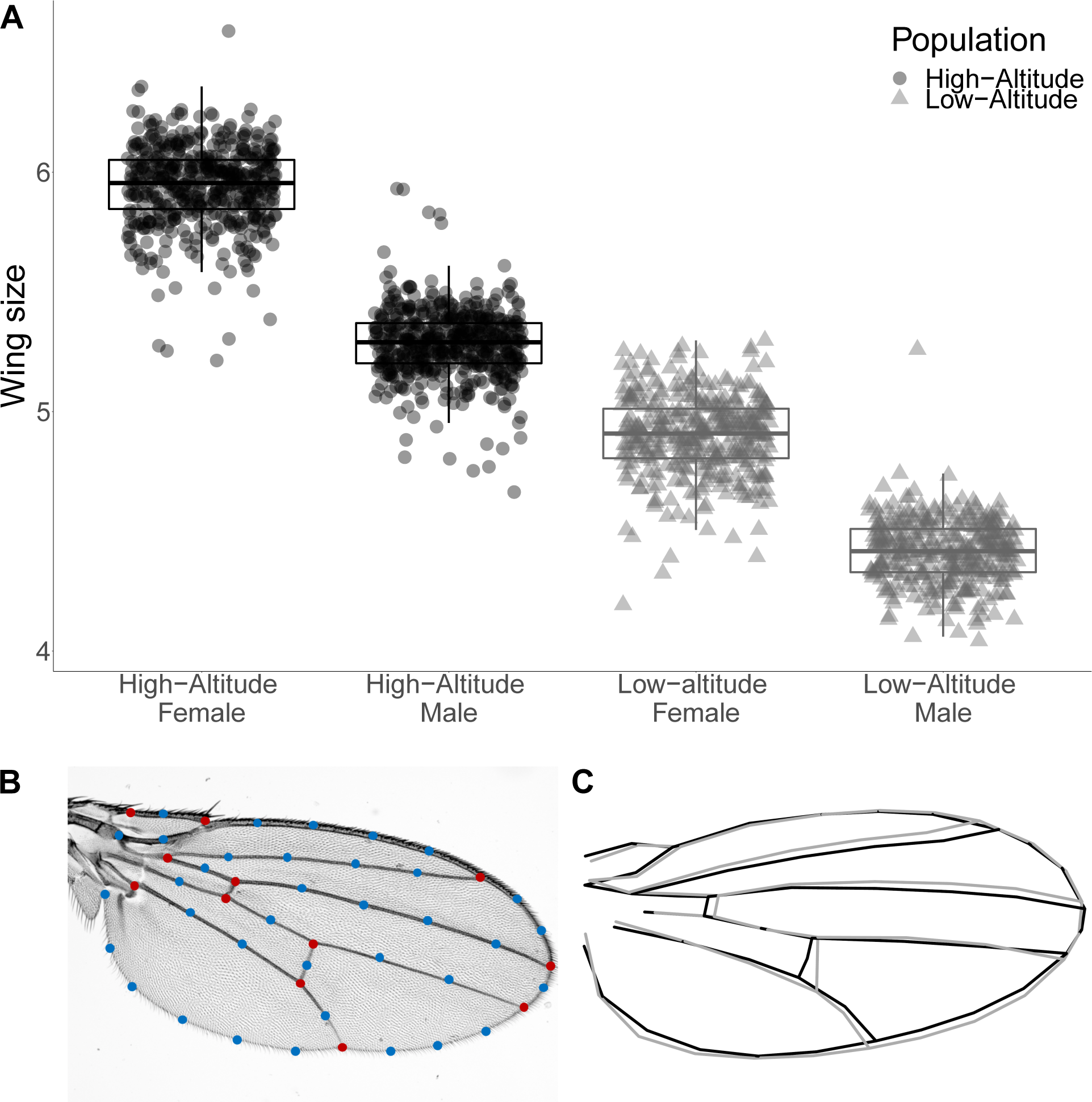
Wing size (centroid size) and shape variation within and among high- and low-altitude populations. (A) High-altitude population has larger wings than the low-altitude population (B) Landmarks (red) and semi-landmarks (blue) used in the analysis of wing shape. (C) Mean difference in wing shape between the high- and low-altitude population, scaled 2x. Procrustes distance between mean shapes of the two populations is 0.013

#### Measuring trichome (cell) density

A subset of strains used for the the initial size and shape analysis were re-imaged with a higher resolution camera. We chose 15 strains from each population as follows: five lines each were chosen with the highest and lowest within-line coefficient of variation (CV) for wing size. Additionally, five lines were chosen at random from each population. This allowed us to maximize the variation we examined within each population. We phenotyped 15-20 males and females from each line. Wings were imaged using an Olympus DP80 camera mounted on an Olympus BX43 microscope, using a 4X objective (total 40X magnification). Images were captured with cellSens Standard (V1.14) software. Cell density was quantified by counting trichomes on the surface of the wing using the ImageJ FijiWings macro (V2.2) (Dobens and Dobens, 2013). Each trichome represents a single cell (Dobzhansky, 1929). We used a 0.0065*mm*^2^ (75×75 px) measurement area in each of 16 different locations in the wing (Figure 4A).

#### Wing size and shape - Macro-environmental canalization

The right wing of each fly was imaged using the same microscope settings as the cell density experiments. Wing size and shape were quantified using the same pipeline as the micro-environmental canalization experiment. We used 3 replicate vials per line per rearing temperature. Strains used for this experiment are outlined in Table S2.

#### Fluctuating Asymmetry of wing size and shape

Left and right wings were phenotyped for two lines from each population (E39 and E73 from high-altitude, Z254 and Z311 from low-altitude, total number of indivudals: 509) to assess fluctuating asymmetry. Duplicate measurements were taken of the left and right wings from 77 individuals chosen randomly from different populations, sexes and rearing temperatures to estimate measurement error. The same phenotyping methods were used as the micro-environmental canalization experiment.

#### Quantification of wing defects

Each wing image was manually scored for venation defects. For the the micro-environmental canalization experiment, proportion of defects was calculated as the ratio of the number of wings with defects to total wings for each line. For the macro-environmental canalization experiment, each individual wing was scored based on whether they have a defect or not, using a binary scale (1 for defect observed, 0 for defect not observed). The proportion of defects for each line was calculated by averaging the scores for all individuals within line and experimental treatment.

### Analysis

Data was analyzed using R (v3.5.1) (R Core Team, 2018) in RStudio (v1.1.456) on a MacBook Pro, running macOS Mojave (V10.14.2). Mixed models were run using *lmer* and *glmer* from the package *lme4* (V1.1.19) (Bates et al., 2015), glmmTMB (Brooks et al., 2017) and *procD.lm* from the package *geomorph* (V3.0.7) (Adams et al., 2018). Generalized linear models were run using *glm* from the *stats* package (V3.5.1) (R Core Team, 2018).

#### Modeling wing size, shape, cell density and wing defects

Linear mixed models were fit with the wing size, and cell density data, and generalized mixed model (binomial distribution, with a logit link) was fit with the wing defects data using population, sex, rearing temperature (for macro-environmental canalization experiment) and their interactions as fixed effects. Where possible, the intercept and sex effects were allowed to vary as random effects of line (strain). Additionally, for the cell density data, wing region was included as a fixed effect to test whether there is variation in cell density across the wing, and an individual level random effect was included to account for the multiple measures per wing.

For the temperature plasticity experiment, we used a model similar to the one described above, but allowing temperature effects to vary according to linear and quadratic effects (using 2nd degree orthogonal polynomials), including interaction effects with temperature and within the random effects of line nested within population.

For wing shape, we fit a multivariate linear model using *procD.lm* estimating the contributing effects of centroid size, population and sex and their interactions as fixed effects, and line nested within population as a random effect. Statistical inference was performed using a randomized residual permutation procedure in geomorph using 1000-2000 permutations for each effect.

#### Estimating among-individual, within-line variation of wing size, shape and cell density

For wing size, among-individual, within-line variation was estimated in two ways for each strain. Using the coefficient of variation 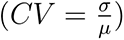 and the median form of Levene’s deviates (used for all formal statistical inference). (Van Valen, 2005; Dworkin, 2005b). For the macro-environmental canalization experiment, sex effects were first modeled out and then CV and Levene’s deviates were calculated for each line at each temperature.

To capture some of the multivariate variational properties for wing shape, we focuses on two measures estimated for each strain. First we used matrix size (total variance), which is the trace of variance covariance matrix for the strain. This is meant to capture overall variation. This is equivalent to the sum of the eigenvalues (Van Valen, 2005). Total variance estimates were multiplied by a factor of 1000.

We also examined trait integration of wing shape using two measures derived from the within-line covariance matrix. Specifically we used matrix eccentricity as well as the standard deviation of its eigenvalues (scaled), (Jones et al., 2003; Kirkpatrick, 2009; Haber, 2011; Pavlicev et al., 2009; Van Valen, 1974). The standard deviation of eigenvalues of the covariance matrix has been used extensively as a proxy for integration (Pavlicev et al., 2009; Cheverud et al., 1983; Haber, 2011). We calculated the relative standard deviation of eigen-values (rSDE) and the relative standard deviation of the eigenvalues scaled by total variance (rSDE2) (Pavlicev et al., 2009; Van Valen, 1974; Haber, 2011). rSDE estimates were multiplied by a factor of 10000 and rSDE2 estimates were multiplied by a factor of 10. The shape of the VCV matrix can also quantified using matrix eccentricity. While typically defined as the ratio between the first two eigenvalues (Jones et al., 2003; Kirkpatrick, 2009), we used a generalization which was the ratio between the largest eigenvalue and the total variance (Haber and Dworkin, 2017), which has been shown to be proportional to rSDE2. Variation due to sex and size was modeled out prior to estimating total variance, eccentricity and rSDE for each strain.

The registration process (Procrustes superimposition) influences covariation within and among landmarks. As such, the use of Procrustes residuals for analysis is of potential concern. However, in our previous study, we demonstrated using a variety of approaches, that at least for *Drosophila* wing shape, results from Procrustes superimposition were extremely similar to those generated via spatial interpolation of the data to generate multivariate variables (Haber and Dworkin, 2017). As such, for this study we used the Procrustes Residuals for simplicity.

CV and Levene’s Deviates, were also calculated for trichome (cell) density for each strain. CV was calculated in two ways. First by averaging the cell density across the wing for each individual and calculating the within-individual CV, and then averaging CV for line. Alternatively, cell density CV was calculated by averaging cell density for each line first and then calculating CV. These two approaches of measuring CV produced similar results and only the first one is used in this paper. Associations between the measures calculated above across strains was performed using a Pearson correlation coefficient.

We examined whether any of the variation measures (CV and Levene’s deviates for size, total variance, eccentricity, rSDE and rSDE2, as well as cell density CV and Levene’s deviates, as response variables) varied due to the effects of population and sex. Generalized mixed models were fit, sex (excluded for for macro-environmental experiment), population, temperature (for macro-environmental canalization experiment) and their interactions as fixed effects. The intercept and where possible sex were allowed to vary as random effects by line (nested within population). Given that all of these responses can only take on continuous positive values we assumed a Gamma distribution with an inverse link function.

#### Fluctuating Asymmetry of wing size and shape

To quantify measurement error, duplicate measures were taken for left and right wing shape for 77 individuals. Wing size measurement error was estimated using an analysis of variance (ANOVA) and wing shape measurement error estimated using a Procrustes ANOVA. For both wing size and shape, individual, side and their interaction were used as effects. The side effect represents directional asymmetry, the individual effect represents variation among individuals and the side:individual interaction term represents fluctuating asymmetry The residual variance in this model estimates measurement error (Klingenberg and McIntyre, 1998; Debat et al., 2009; Palmer and Strobeck, 1986, 2003; Palmer, 1994). We compared the variation of the side:individual interaction term with the residual variation in order to determine whether the measurement error was negligible with respect to fluctuating asymmetry (Supplementary Tables S24 and S25).

Fluctuating asymmetry of wing size was calculated using standard FA indices: FA1 (*FA*1 = *|R − L|*) and 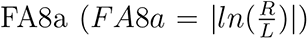 (Palmer and Strobeck, 1986, 2003; Palmer, 1994).

In order to estimate and assess differences in developmental stability (based on FA), we fit a generalized linear mixed model using the FA indices for wing size *FA*1 and *FA*8*a* as response variables, temperature, sex, population and their interactions as fixed effects. Random effects of the intercept, sex and temperature were allowed to vary according to line nested within population. A Gamma distribution and an inverse link function for the response were used. The FA component for shape was extracted for each specimen using the *bilat.symmetry* function from *geomorph* to remove directional asymmetry. As a confirmation of this analysis, wing shape FA was calculated as the Procrustes distance between the left and right wing for each individual *P D_RL_*. We fit a generalized linear model using *P D_RL_* as response variables, temperature, sex, population and their interactions as fixed effects and line as random effect assuming a Gamma distribution and an inverse link function. As a confirmation of the FA analysis, morphological disparity analysis was performed to compare the difference in FA among groups. Each analysis provided largely similar results and only the first two are shown.

## Results

### Wing size, shape and wing defects vary between high- and low-altitude populations

We first confirmed differences in trait means across populations. Consistent with previous findings (Lack et al., 2016a; Pitchers et al., 2013; Fabian et al., 2015), the high-altitude population has substantially larger wing size compared to the low-altitude population (Figure 1A, Table S4). Wing shape also varies in a manner consistent with previous studies (mean Procrustes distance of 0.013 between populations) (Pitchers et al., 2013) shown in Figure 1C and Table S5.

Consistent with Lack et al. (2016a), flies from the high-altitude population have a greater proportion of wing defects compared to the low-altitude population (Figure S1, Table S8). On average, 26.7% (CI 22.8% - 29.9%) of high-altitude females and 22.0% (CI 18.6% - 25.8%) of males show such defects. In contrast, the average for the low-altitude females is 10.6% (CI 7.53% - 14.6%) and 12.2% (CIs 8.22% - 17.6%) for males.

#### Micro-environmental variation for wing size is similar between high- and low-altitude populations

While it was previously demonstrated that the high altitude population is genetically decanalized (Lack et al., 2016a), it is unclear whether this is also associated with any form of environmental decanalization. To enable comparisons with previous studies we used both the coefficient of variation (CV) and Levene’s deviates to measure among individual, within-line variation. CV was plotted for ease of interpretation, but all statistical analyses were performed using Levene’s deviates (Van Valen, 2005; Dworkin, 2005b). However, Levene’s deviates and CV are highly correlated (high-altitude r = 0.89 CIs 0.81 - 0.94; low-altitude r = 0.98 CIs 0.94 - 0.99) (Figure S3B). As shown in Figure 2A, Figure S3A and Tables S6 and S7, measures of among individual, within-strain variability are similar between the high-altitude and low-altitude populations.

**Figure 2:**
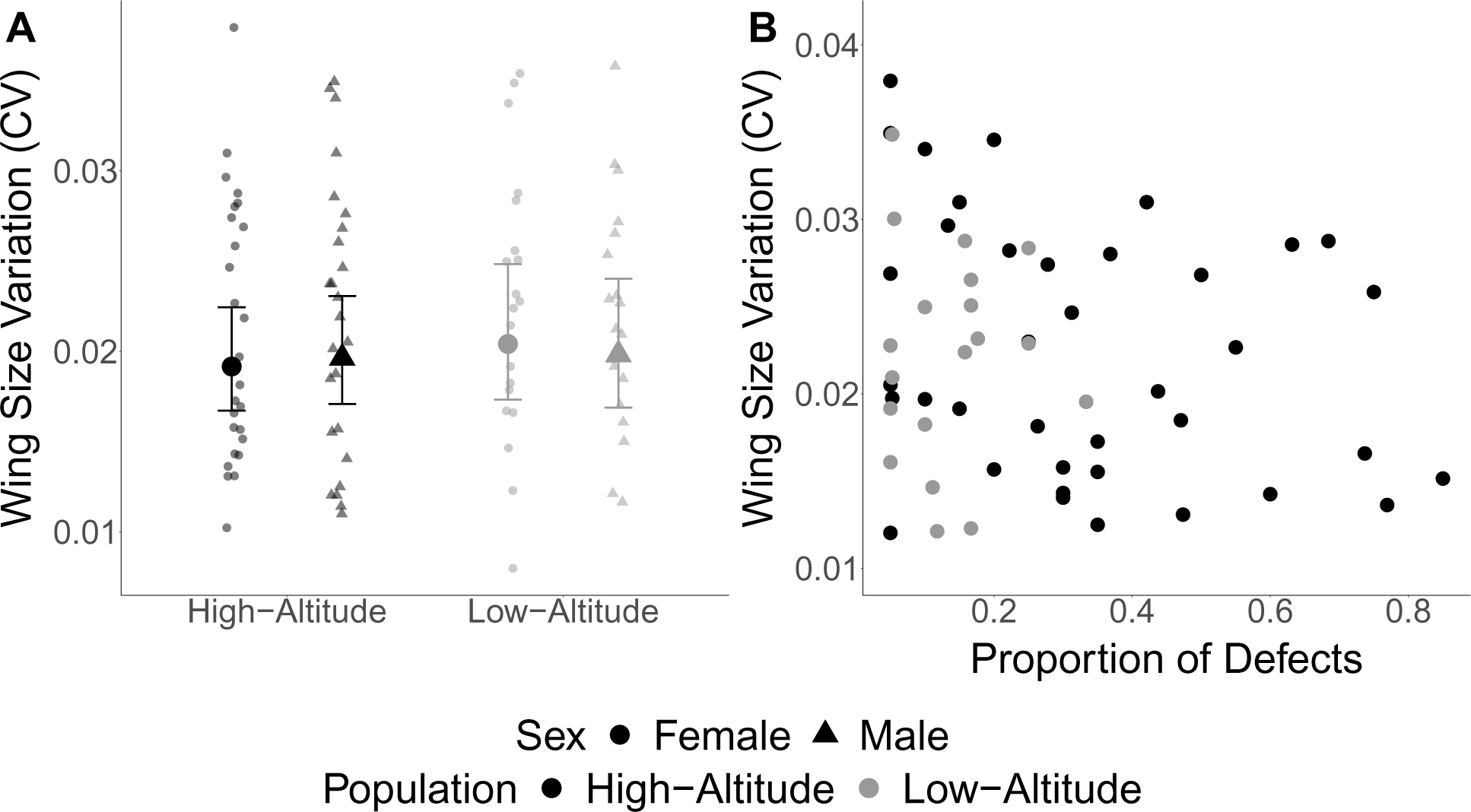
Within-line, among individual variation for wing size is similar across high- and low-altitude populations. (A) Within-line coefficient of variation for wing size is similar in high- and low-altitude populations. Large symbols represent fitted values, small symbols represent coefficient of variation by line. Error bars are 95% CI (B)Within-line variation for wing size is not correlated with proportion of defects in both the high-altitude population *r* = 0.28 (CIs 0.55 - 0.045) or the low-altitude population *r* = 7.45 10*^−^*^3^ (CIs 0.46 - 0.45)

We examined the relationship between CV and proportion of defects, which showed a weak negative correlation, in the high-altitude population *r* = *−*0.28 (CIs *−*0.55 - 0.045), and a correlation close to zero in the low-altitude population *r* = *−*7.45 *×* 10*^−^*^3^ (CIs *−*0.46 - 0.45) (Figure 2B, Figure S4A) although confidence intervals included zero for both populations. Similar results were observed when comparing within-line Levene’s Deviates with proportion of defects (Figure S4A).

### Micro-environmental variation for wing shape is similar between high- and low-altitude populations

Strains derived from high- and low-altitude populations have similar levels of wing shape variation measured as the total variance (matrix size). This is also true for measures of integration (Figure 3 and Tables S9 and S10). We observed this using both the relative standard deviation of eigenvalues and eccentricity of the covariance matrices (Figure S6). Similar to the patterns for wing shape among populations, there is little evidence that total variance, eccentricity, rSDE and rSDE2 are correlated with frequency of wing defects (Figures S2 and S5) in either high-altitude (Total variance: r = 0.16 CI *−*0.26 - 0.53; eccentricity: r = *−*0.24 CI *−*0.59 - 0.18; rSDE: r = 7.6 *×* 10*^−^*^3^ 95% CI *−*0.40 - 0.41; rSDE2: r = *−*0.29 95% CI *−*0.62 - 0.12) or low-altitude populations (total variance: r = 0.34 CI *−*0.24 - 0.74; eccentricity: r = *−*0.33 CI *−*0.73 - 0.24; rSDE: r = 0.12 95% CI *−*0.44 - 0.61; rSDE2: r = *−*0.17 95% CI *−*0.65 - 0.39).

**Figure 3:**
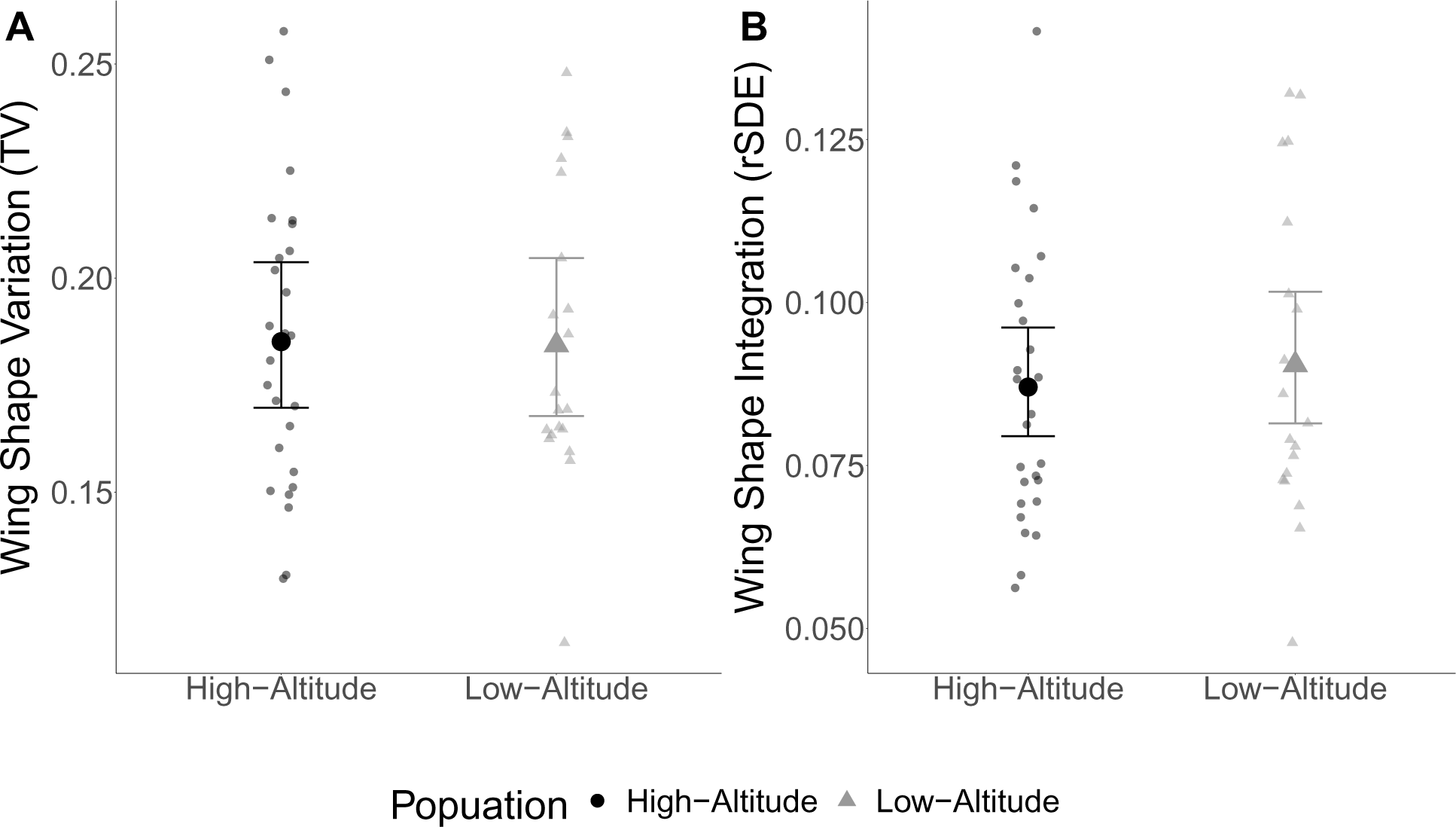
Similar patterns of within-line measures of variability for wing shape between high- and low-altitude populations using (A) a measure of wing shape variation, total variance of the VCV matrix (values multiplied by 1000), and (B) measure of wing shape integration using relative standard deviation of the eigenvalues of the VCV matrix (rSDE; values multiplied by 10000). Error bars are 95% CIs

### Cell Density varies across the wing, between population and sexes

Consistent with previous work, we observed that average cell density is lower (cell size is greater) in the high-altitude population relative to the low-altitude population (Figure 4B) (Fabian et al., 2015; Lack et al., 2016b) as well as for females versus males (Alpatov, 1930; Dobzhansky, 1929). Most studies count trichomes (cells) in a single small region of the wing. Yet, cell sizes are known to vary in different regions of the wing from less than 7.9*µm* to greater than 11.3*µm* in diameter (González-Gaitán et al., 1994). To account for local effects, we measured cell density in 16 regions across the wing (Figure 4A, Figure S7). While cell density varies considerably across the wing, with some intriguing interactions between sex, region of the wing and population, the overall pattern as observed in previous work (Lack et al., 2016b; Fabian et al., 2015) remains (Figure S7; Table S11).

**Figure 4:**
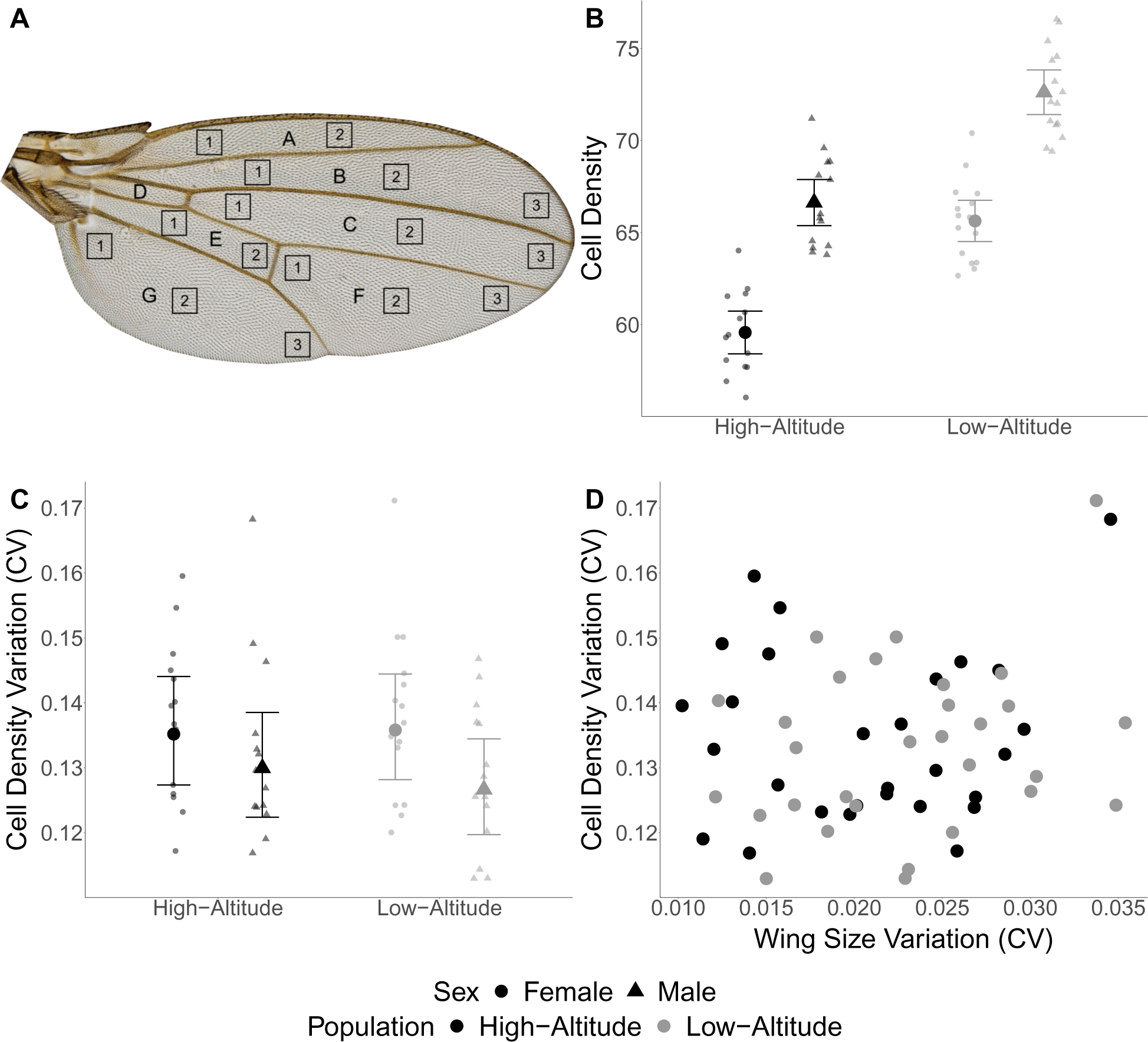
Mean cell density show population and sex differences, but no changes in variability. (A) Wing regions used for cell density measurement. Squares represent a 0.0065*mm*^2^ measurement area. (B) Cell density varies between the sexes and between the high- and low-altitude populations. Error bars are 95% CIs (C) Among-individual, within-line variation measured as CV for cell density is similar between high- and low-altitude populations. (D) Within-line Cell density variation and within-line wing size variation (CV) are not strongly correlated in either the high-altitude population (*r* = 0.076 CI *−*0.31 - 0.44), and the low-altitude population (*r* = 0.25 CI *−*0.12 - 0.56)

### Variation within and among individuals for cell density is not associated with among individual variability in size or shape

After we confirmed and expanded upon the previously demonstrated association between cell density and wing size with respect to population and sex, we asked whether variation in cell density, within the wing was directly associated with variation among individuals in wing size and shape. That is, do lines that show the greatest degree of among individual, within-line variance for wing size and shape also show the greatest variance for cell density within and between individuals? We calculated the within-line CV for cell density across the wing in order to determine if there are any differences in within-line variation for cell density between the high- and low-altitude populations. As shown in Figure 4, we did not observe substantial differences in cell density CV between populations but observed an effect of sex that was consistent across both populations (Figure 4C; Table S12). Similarly, we did not observe differences in within-line cell density Levene’s deviates between the high- and low-altitude populations (Figure S8A; Table S13).

Additionally, we did not observe a strong association between within-line cell density CV and within-line wing size CV for either the high-altitude population (*r* = 0.076 Cis *−*0.31 - 0.44) or the low-altitude population (*r* = 0.25 CIs *−*0.12 - 0.56)(Figure 4D). We observed a weak negative correlation between within-line cell density CV and within-line total variance,in the low-altitude population (*r* = *−*0.41 CIs *−*0.67 - *−*0.060;), but we did not observe any correlation between within-line cell density CV and within-line total variance for the high-altitude population (high-altitude *r* = 0.062 CIs *−*0.32 - 0.42). Similarly, we did not observe any association between within-line cell density CV and within-line eccentricity in either the high- or the low-altitude population (high-altitude *r* = 0.28 CIs *−*0.10 - 0.59, low-altitude *r* = *−*0.30 CIs *−*0.60 - 0.063) (Figure S8 C, D). Further, we did not observe any evidence for associations between within-line cell density CV and proportion of wing defects for both the high- and low-altitude populations(high-altitude *r* = *−*0.043 CIs *−*0.44 - 0.37, low-altitude *r* = 0.36 CIs *−*0.076 - 0.68) (Figure S8B)

### Temperature induced plasticity

To assess whether patterns of phenotypic plasticity varied among populations (macro-environmental canalization) we reared strains derived from both populations at three temperatures. Consistent with previous studies, wing size is larger for flies raised at lower temperature and the reaction norms showed modest evidence of non-linearity (David et al., 1994; Partridge et al., 1994; James et al., 1997). Our data suggest that mean wing size of the high-altitude population may be more plastic compared to the low-altitude population (Figure 5A, Table S14, Table S15). We observed an increase in the Procrustes distance between mean shapes of the high- and low-altitude populations as temperature increases (Figure S11).

**Figure 5:**
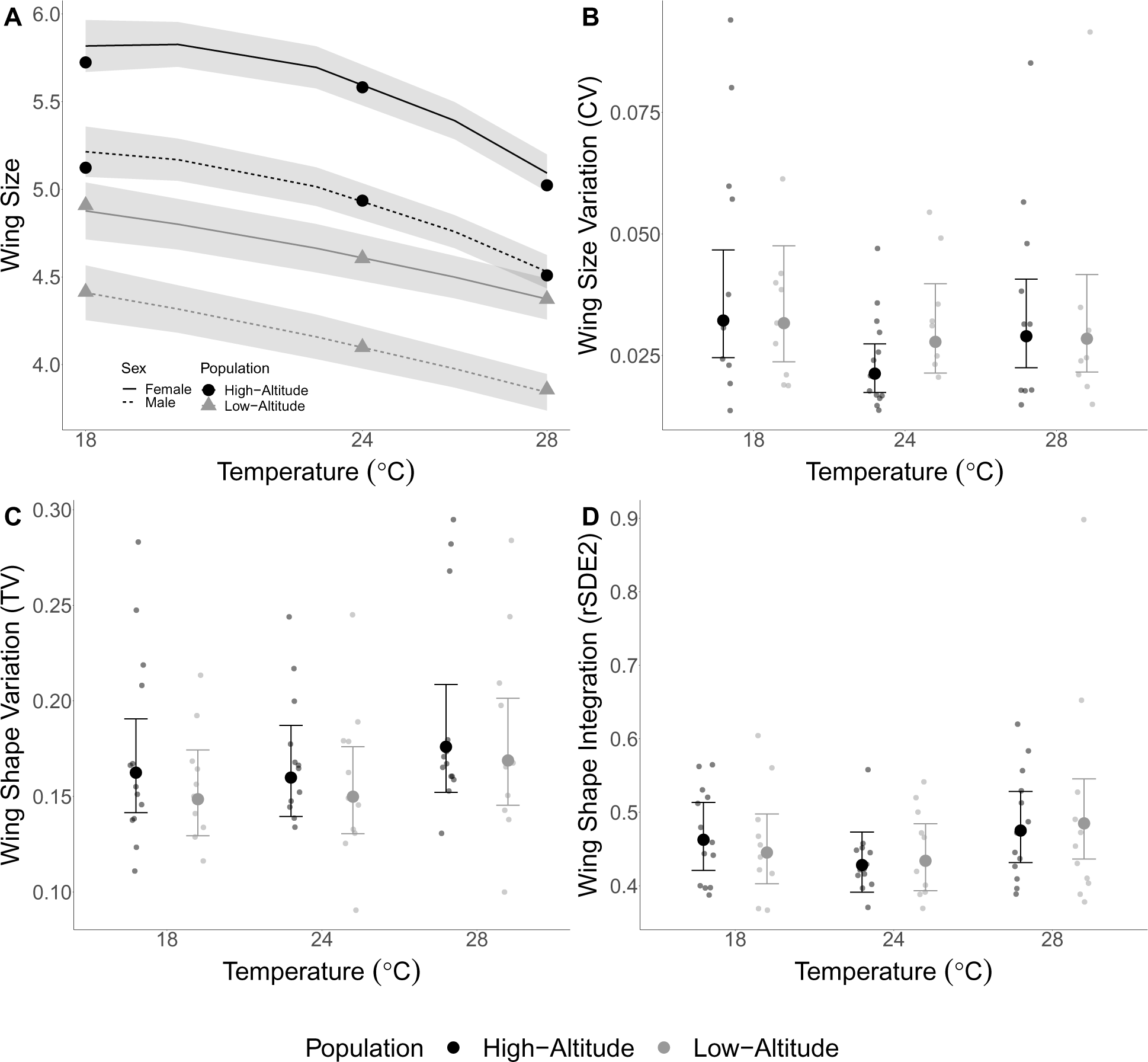
Considerable plasticity for mean wing size under different temperature rearing environments, but with minimal changes in trait variation (A) Mean changes in wing size (centroid size) (B) Within-line variation for wing size is similar across sexes and populations within each temperature treatment. (C) Within-line variation for wing shape measured as the total variance of the VCV matrix (multiplied by a factor of 1000) and (D) Within-line wing shape integration measured as rSDE2 (relative standard deviation of the eigenvalues scaled by the total variance) are similar between the high- and low-altitude populations across the different rearing temperatures. Gray shading and error bars are 95% CIs

We quantified the proportion of wing defects for the high- and low-altitude populations at the three different rearing temperatures. Consistent with our previous results, the high-altitude population has a greater proportion of defects than the low-altitude population, however, we did not observe substantial differences in proportion of defects due to temperature (Figure S9A, Table S20).

In general, we did not observe any differences in the high- and low-altitude populations for among individual, within-line measures of variability at each temperature treatment. For the within-line CV for wing size, we observed an increase at both 18°and 28°C for high-altitude females and an increase in CV at 18°but not at 28°C for high-altitude males. Within-line CV for low-altitude males and females is consistent across temperatures (Figure 5B; Table S16). We observed a similar pattern when using Levene’s deviates as we did for CV (Figure S9B; Table S17), with a modest effect of rearing temperature but little evidence for population level differences. For within-line wing shape total variance, we observe a consistent increase with temperature for both high- and low-altitude populations (Figure 5C; Table S18). Degree of integration of wing shape is similar across populations and temperatures (Figure 5D; Table S19). This pattern holds whether examining rSDE or eccentricity of the covariance matrix (Figure S9C, D).

### Fluctuating asymmetry for wing size and shape is similar between high- and low-altitude populations

While among-individual, within-genotype variation and within-individual (among-sides) variation might be expected to capture similar aspects of developmental stability, empirical work shows that these measures do not always agree with correlations of *∼ between*0.07*and*0.6 for wing size and *∼* 0.35 *−* 0.48 for shape (Debat et al., 2006, 2009; Breuker et al., 2006). Thus we measured asymmetry for both wing size and shape in a subset high- and low-altitude lines (2 strains for each population). We first estimated measurement error for wing size and shape and determined that measurement error was negligible with respect to FA for size (Table S25) although had a larger impact on shape (Table S26). Using FA8 as an index for wing size FA, we compared developmental instability for high- and low-altitude populations at three temperatures (18°C, 24°C and 28°C). We did not see clear evidence of differences in FA8 between high- and low-altitude populations or across the different rearing temperatures, except for consistently lower FA8 in the high-altitude males at all temperatures (Figure 6A; Table S22). For comparison we also report FA1 as a measure of developmental instability ((Figure S12; Table S21)), but importantly FA1 does not account for mean trait size and should be interpreted with caution.

**Figure 6:**
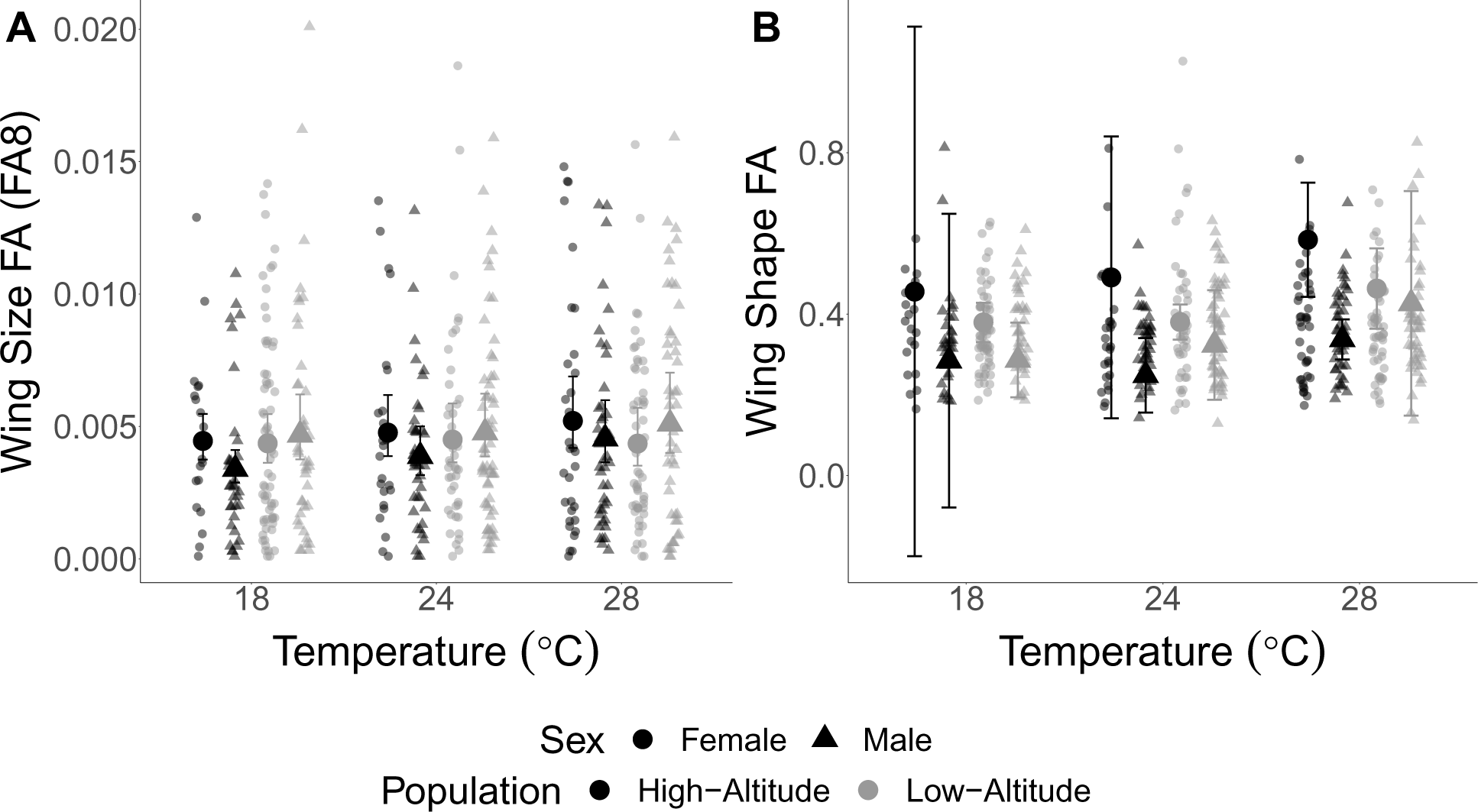
Inconsistent differences in fluctuating asymmetry among the high- and low-altitude populations for (A) wing size fluctuating asymmetry measured as *FA*8 and (B) wing shape fluctuating asymmetry between the high- and low-altitude populations across temperatures. High-altitude males have consistently lower fluctuating asymmetry and high-altitude females have consistently greater wing shape fluctuating asymmetry across temperatures. Large symbols represent population means, small symbols represent individuals, error bars represent 95% CIs

We measured FA for wing shape in two different ways. First, we calculated FA by removing directional asymmetry (DA) and then calculating the Procrustes distance (PD) for the FA component for each individual. The high-altitude females had a slight, but consistent increase in FA across all three temperatures while the other groups had similar FA to each other and across the different temperatures (Figure 6B Table S24). We also calculated Procrustes Distance between left and right wings (*P D_LR_*). We observed a similar pattern as we did using to first method to calculate wing shape FA, where there is consistently greater *P D_LR_* in the high-altitude females across temperatures (Figure S12B; Table S23).

## Discussion

Since the proposal of the congruence hypothesis, researchers have tested for associations between genetic and environmental canalization. To date, empirical evidence has been equivocal. The majority of studies that support the congruence hypothesis were conducted in RNA viruses and microRNAs *in vivo* and *in silico*, which may not necessarily representative of multi-cellular organisms (Szöllősi and Derényi, 2009). Studies in other systems have not provided evidence to support the congruence hypothesis (Dworkin, 2005a,c). However, most empirical studies were conducted using lab domesticated lineages, laboratory-induced mutations, and with arbitrary measures of genetic canalization which may not be representative of naturally occurring phenomena that would lead to congruent evolution of genetic and environmental canalization.

In this study, we address these issues using a naturally occurring system across adaptively diverged populations. The high-altitude population was previously demonstrated to have reduced mutational robustness (via mutagenesis), and these effects were pleiotropically linked to variants influencing changes in wing and body size that appear to be targets of selection (Lack et al., 2016a). Using strains derived from both high- and low-altitude populations, we examined multiple measures of environmental canalization. Despite recapitulating previously observed divergence in wing size & shape (Pitchers et al., 2013; Lack et al., 2016a,b) (Figures 1, 5A, S11), cell size (Figure 4, Figure S7) and frequencies of mutational defects (Figure 2B, Figure S1), we did not observe any evidence for associations between genetic and micro-environmental canalization (Figures 2, 3, 4C-D, 5B-D). Additionally, measures of among individual, within-line variance for wing size and shape were not correlated with the proportion of mutational defects both the high- and low-altitude populations (Figures 2B, 4D, Figures S2, S4, S8B, S10). We did observe greater temperature induced plasticity of mean wing size in the high-altitude population (Figure 5A). We observed a subtle increase in within-line variation for wing size at 18°C and 28°C compared to 24°C in both the high- and low-altitude population for wing size, and this increase was greater in the high-altitude population (Figure 5B, Figure S9B), although there is at best, marginal evidence for a significant treatment effect of population or its interaction with rearing temperature (Table S16). Intriguingly, we observed a decrease in developmental stability for wing size (measured using fluctuating asymmetry) for high-altitude females across all temperatures (Figure 6). Yet, we did not observe this same pattern in the males derived from the same population, nor did we see any increase in qualitative wing defects at varying temperatures and the measures of among-individual, within-line of variation were not correlated with the proportion of defects at any of the developmental temperatures (S10). Therefore, our results are largely inconsistent with congruent evolution of genetic and environmental canalization and that with respect to adaptation to life at high altitudes driving changes in both trait means and variances, they are likely to evolve via separate underlying mechanisms.

Our study is one of the few to test of the congruence hypothesis using strains derived from natural populations with known evolutionary histories. However, we are aware of several important caveats. Our study compares a single high-altitude population to a single low-altitude population. Replication of our experiments with additional populations from independent altitudinal clines would provide stronger support of our findings. The strains used in this study were collected approximately 2 years prior to the first experiment and seven years prior to the temperature manipulation experiment. As such both drift (due to the initial inbreeding process) some degree of lab adaptation may have occurred. As we used multiple strains from each population, the impact of drift with respect to allele frequencies should be modest. Additionally lab domestication should be weak as *N_e_* is extremely small within each strain. Although the two populations have modest genetic differentiation (*F_ST_* = 0.15), lines derived from African populations tend to have have high residual heterozygosity even after 8 generations of inbreeding (Lack et al., 2016a). However, given that our results for mean size and shape of the wings and cell densities recapitulate previous findings, the impact of both drift and lab domestication appear to be minor.

While the variants contributing to divergence in size both between the high- and low-altitude population (Lack et al., 2016a) and under artificial selection derived from the low-altitude population (Groth et al., 2018) appear to be pleiotropically linked to the mechanism influencing sensitivity to mutational perturbation, these are not in fact the same traits. The penetrance of wing abnormalities among lines derived from high altitude, and the increased sensitivity under mutagenesis may reflect one aspect of genetic canalization (that is linked to variants influencing mean size and shape), but they do not necessarily influence variance for these traits. Indeed, under high temperature stress (31°C) one of three replicates of lineages artificially selected for increased size (from a low-altitude ancestral population) showed a substantial increase in penetrance of wing abnormalities (Groth et al., 2018). Interestingly we observed no increase in such abnormalities at our high temperature rearing (28°C) for the high-altitude strains. Whether this reflects insufficient stress or a difference in response is unclear. However, it is clear that the degree of genetic correlation between trait means and variances for wing size, shape and penetrance of abnormalities is complex.

While the work of Lack et al. (2016a,b) and Groth et al. (2018) clearly demonstrate evolutionary changes in genetic canalization associated with adaptive trait evolution (body size and wing morphology), the most likely evolutionary scenario is one of the loss of genetic canalization associated with the pleiotropic effects of variants that have contributed to the evolution of mean size at high altitudes. This is reminiscent of the evolution of insecticide resistance in blowflies which showed a pleiotropic increase in fluctuating asymmetry due to the resistance locus or in sticklebacks with the effects of the *Eda* locus on both the expression of armour plates, and on fluctuating asymmetry of plates (McKenzie and Clarke, 1988; Morris et al., 2019). Consistent with Waddington’s model for the evolution of canalization, it could be that modifiers that increase the mutational robustness of wing morphology in *Drosophila* have not yet risen to appreciable frequency in the high altitude population, as has occurred with the modifier variants influencing asymmetry in the blowflies (Davies et al., 1996). Indeed it is not yet clear whether wing size is near its optima for the high altitude population, and whether that is necessary for the evolution of canalization.

Based on what is known about the genetic architecture of body size, wing size and wing shape the fact that genetic de-canalization occurred is surprising. The mutational target size of body size, wing size and wing shape are quite large (Weber et al., 2005; Houle and Fierst, 2012; Carreira et al., 2008, 2011). Similarly these traits harbour extensive standing genetic variation in populations and are polygenic in nature (Weber, 1990b,a; Weber et al., 1999; Mezey et al., 2005; Mezey and Houle, 2005). As such the expected modest changes in individual allele frequencies would seem to be unlikely to result in changes in canalization. Yet that is exactly what has been observed in the Ethiopian high altitude population which increased its wing (and body) size, and also from the low altitude population artificially selected for large wing size. Body and wing size has been a frequent subject of study in *Drosophila*, but until now this pattern has not been previously observed. As such future work both examining additional populations that vary for size and also on on identifying variants influencing the adaptive divergence in wing size and morphology and how they shape mutational robustness are necessary.

Where does this leave the congruence scenario? While the results from the current study, and some previous studies (Dworkin, 2005a,c; Borenstein and Ruppin, 2006) are not consistent with the congruence hypothesis, it is perhaps best to consider under what conditions the direct evolution of genetic canalization or its evolution as a correlated response are probable. The model observation that finds that deleterious alleles are purged efficiently enough that adaptive genetic canalization will not have the opportunity to evolve does not consider that deleterious alleles are not always purged efficiently for numerous reasons (pleiotropy, linkage with beneficial mutations, drift, GxE, fluctuating selection etc). Indeed, with the system we investigated in this study, the reduction in mutational robustness appears to be a direct pleiotropic consequence of the allelic effects on organismal size (Lack et al., 2016a; Groth et al., 2018). In such instances, genetic canalization may evolve to suppress deleterious mutational effects (ie. alter patterns of pleiotropy). As has been demonstrated previously, the likelihood of evolving either genetic or environmental canalization is in part a function of the fitness load imposed by the frequency and magnitude of environmental and genetic perturbations (Proulx and Phillips, 2005; Hermisson et al., 2003). However, our knowledge of the distribution of this fitness load and the frequency of relevant environmental perturbations is limited. While studies of mutational load give us some idea of the distribution of fitness effects of new mutations, this is less clear in natural environments. Indeed, it has been argued that new allelic combinations produced due to the normal processes of mating and recombination may act as a “genetic perturbation” to the input of new mutations (Stearns et al., 1995), as genetic backgrounds are constantly shuffled. Alternatively, it may be that genetic canalization may be beneficial when it occurs, but rarely the result of persistent and direct selection (Wagner et al., 1997; Gibson and Wagner, 2000; Visser et al., 2003; Siegal and Bergman, 2002; Proulx and Phillips, 2005). Experimental evolution and artificial selection may continue to be the strongest framework to test the theory and understand under what conditions both genetic and environmental canalization are direct targets of selection. This approach should be coupled with studies of adaptively diverged natural populations that are likely to share the appropriate evolutionary history to address these questions (i.e. (Morris et al., 2019; McKenzie and Clarke, 1988)). Finally, this work suggests that trying to clearly delineate between selection for ‘environmental’ and ‘genetic’ canalization may be difficult given the interplay between genotypic and environmental effects in terms of trait expression and variation.

## Acknowledgements

We wish to thank Dr. John Pool and Dr. Justin Lack for sending strains. We thank Megan Cermak, Christian Marier, Sachin Davis, Melissa Rezik, Ravina Dhami, and Vikram Bhagavat for help with dissections and imaging. Dr. Greg Gibson and Dr. John Pool provided feedback on an earlier version of this manuscript. Dr. Michael Collyer and 2 anonymous reviewers provided helpful feedback that improved the clarity of this manuscript. This work was funded by an NSERC (Canada) Discovery and Discovery accelerator grant to ID.

## Supplemental Figures

**Figure S1:**
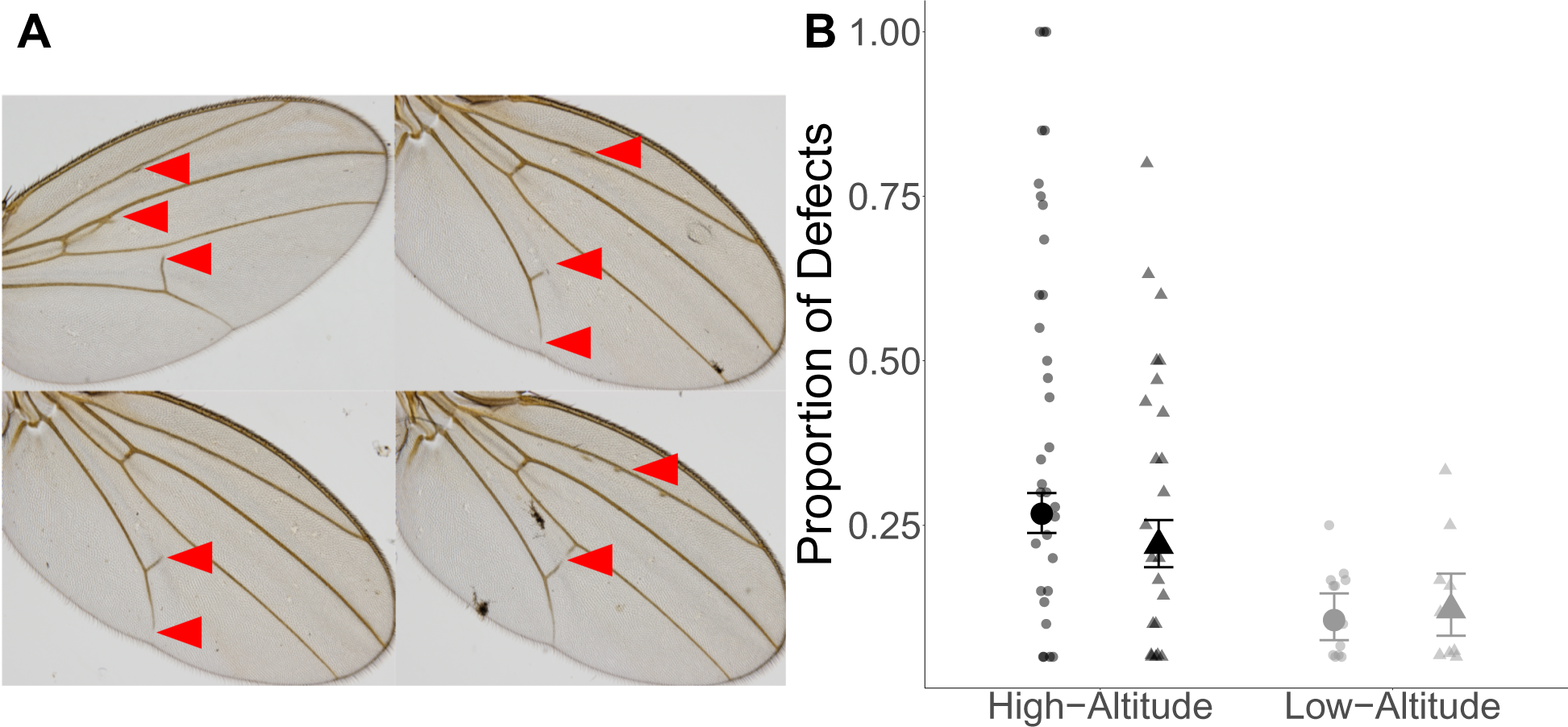
High-altitude population has a higher frequency of wing defects compared to the low-altitude population. (A) Examples of wing defects seen in the high-altitude population (red arrowheads pointing to venation defects). Venation defects represented include additional longitudinal veins (top left) or small pieces of vein material (top left and top and bottom right), incomplete posterior cross vein (all 4 wings) and incomplete L5 vein (bottom left) (B) Proportion of wing venation defects within lines is much greater in the high-altitude population than the low-altitude population. Circles - females, triangles- males.

**Figure S2:**
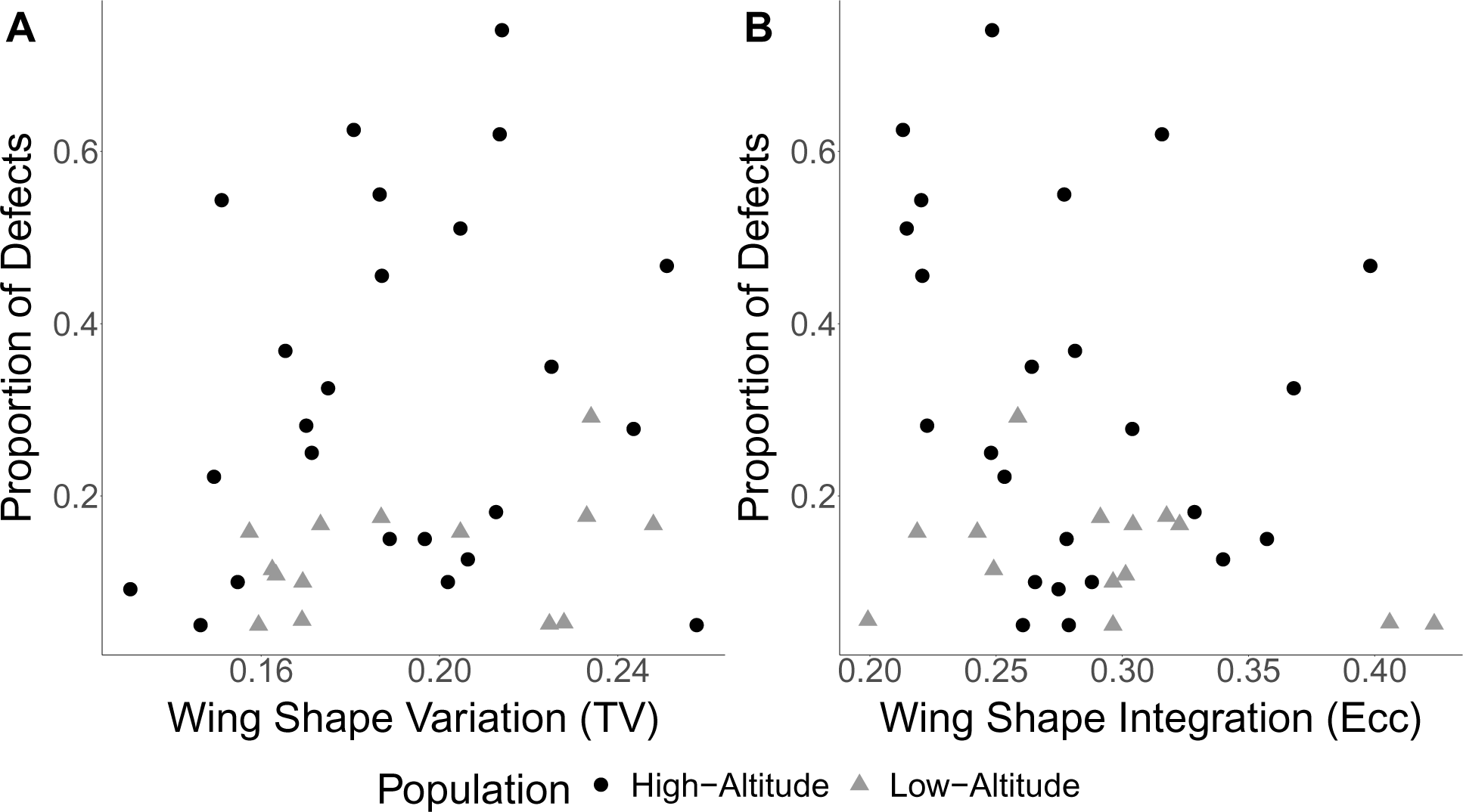
Within-line measures of variability for wing shape are not correlated with proportion of wing defects (TV - Total variance: r = 0.16 95% CI *−*0.26 - 0.53;Ecc - Eccentricity: r = *−*0.24 95% CI *−*0.59 - 0.18) or the low-altitude population (Total variance: r = 0.34 95% CI *−*0.24 - 0.74; eccentricity: r = *−*0.33 95% CI *−*0.73 - 0.24)

**Figure S3:**
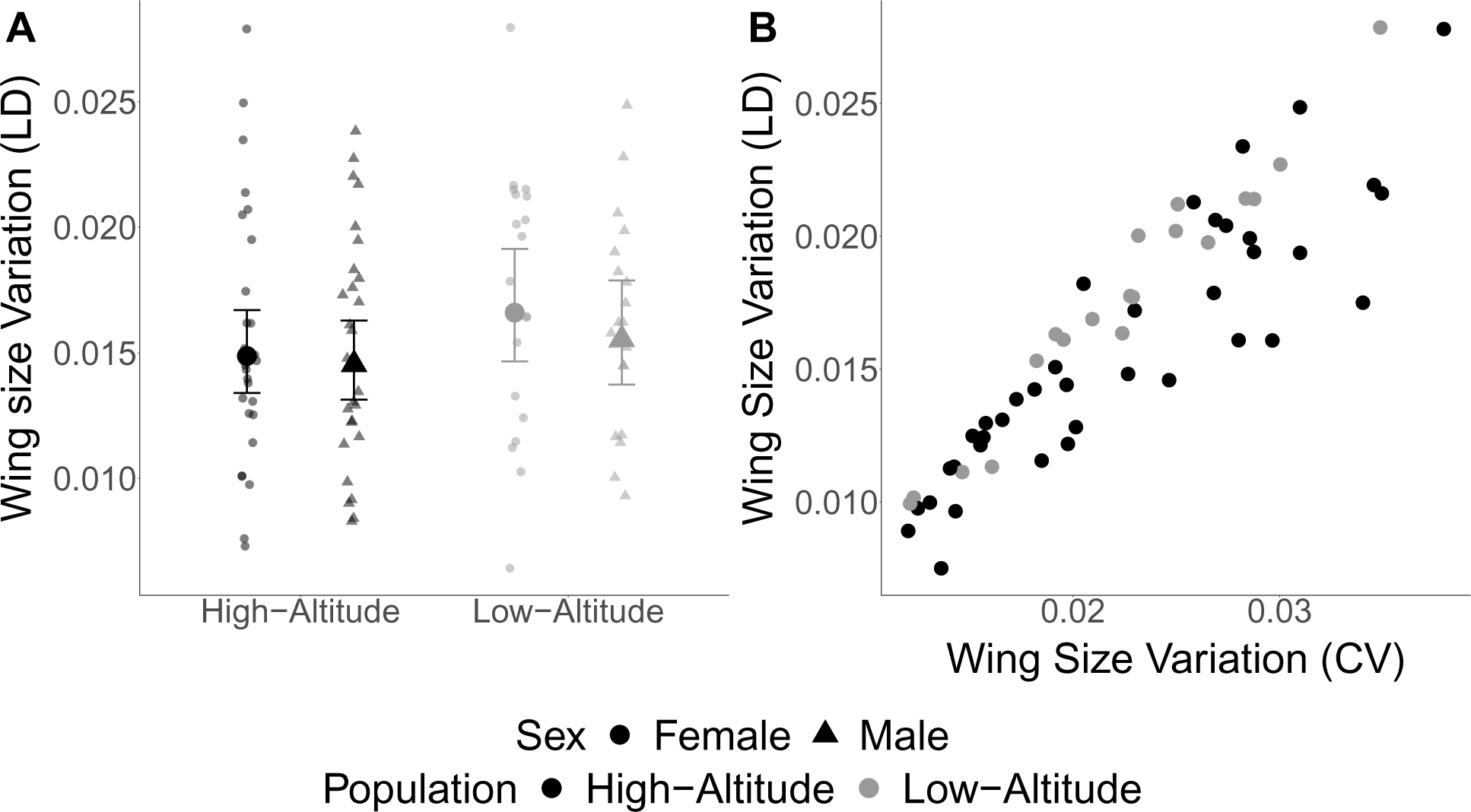
Within-line wing size variation measured as Levene’s Deviates. (A) Consistent with the results using the coefficient of variation, Levene’s deviates are similar for the high- and low-altitude populations. (B) The two measures of within-line variation, Levene’s Deviates and CV, are highly correlated (high-altitude r = 0.89 95% CIs 0.80 - 0.94; low-altitude r = 0.98 95% CIs 0.94 - 0.99)

**Figure S4:**
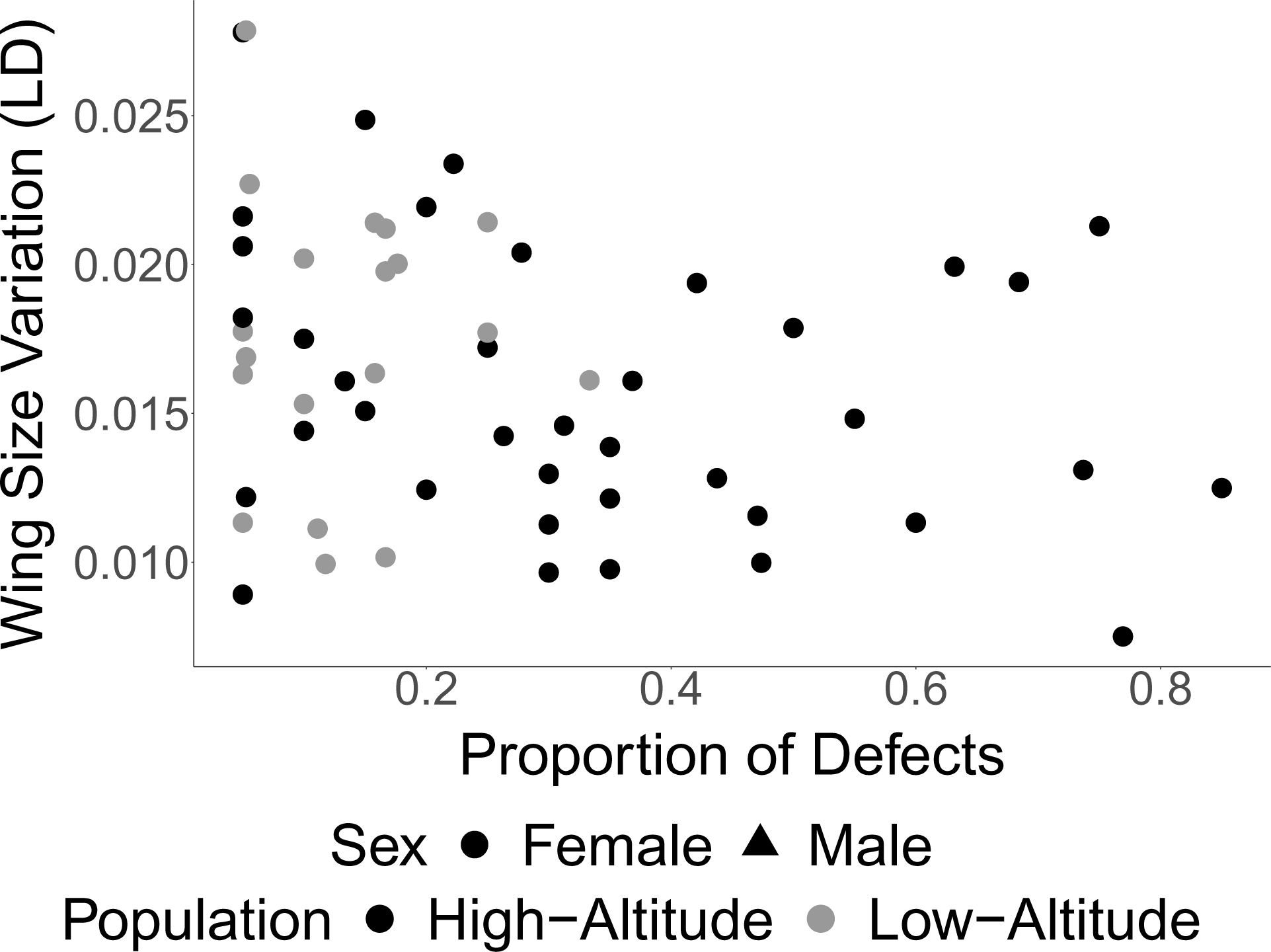
Within-line variation for wing size measured as Levene’s deviates are not correlated with within-line proportion of defects in either the high-altitude population(r = *−*0.27 95% CIs *−*0.54 - 0.055) and the low-altitude population (r = *−*0.0032 95% CIs *−*0.46 - 0.45).

**Figure S5:**
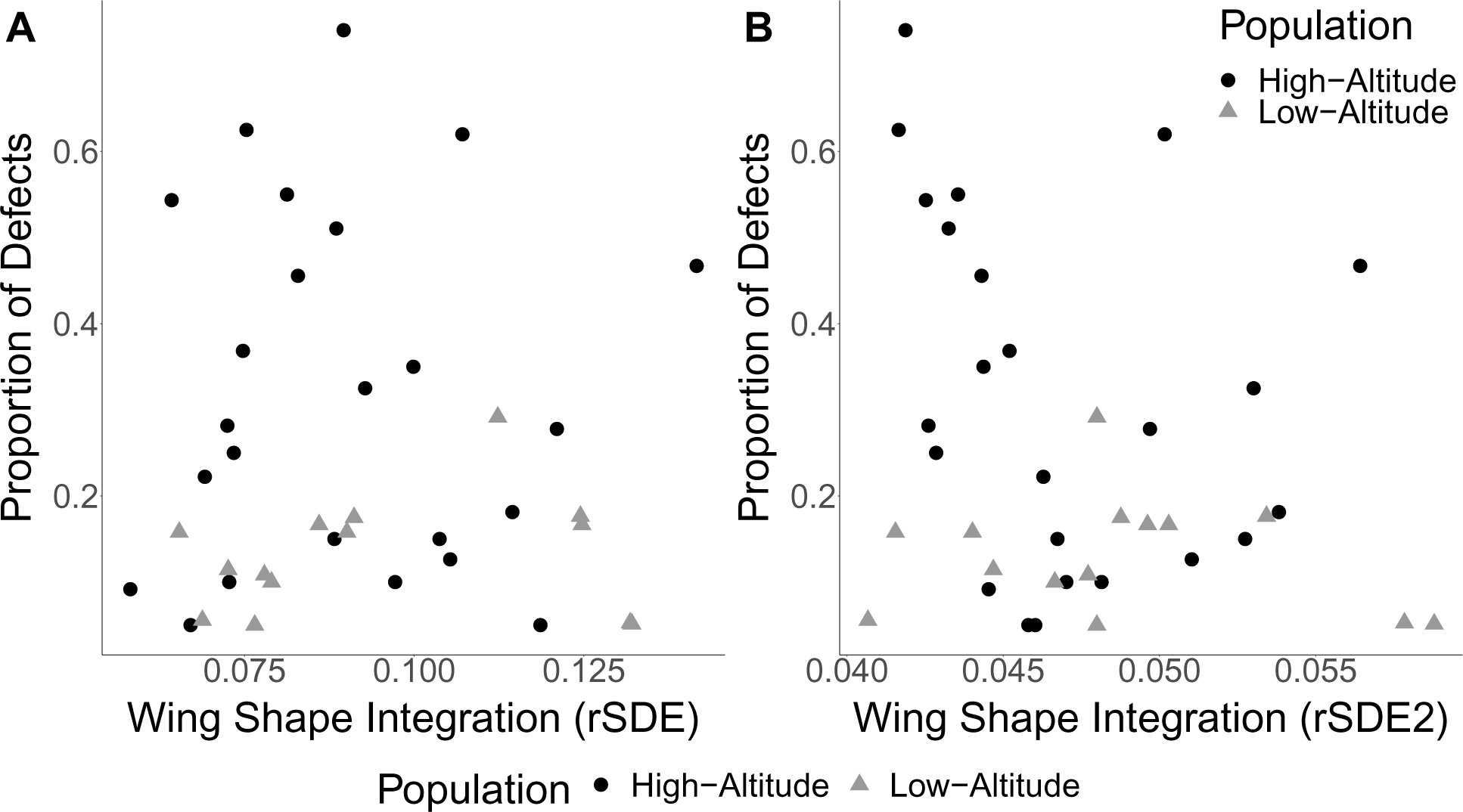
Within-line measures of variability for wing shape measured as wing shape integration using the relative standard deviation of the eigenvalues (rSDE - multiplied by 10000 and rSDE2) are not correlated with proportion of wing defects (rSDE: r = 7.6 *×* 10*^−^*^3^ 95% CI *−*0.40 - 0.41; rSDE2: r = *−*0.29 95% CI *−*0.62 - 0.12) or the low-altitude population (rSDE: r = 0.12 95% CI *−*0.44 - 0.61; rSDE2: r = *−*0.17 95% CI *−*0.65 - 0.39)

**Figure S6:**
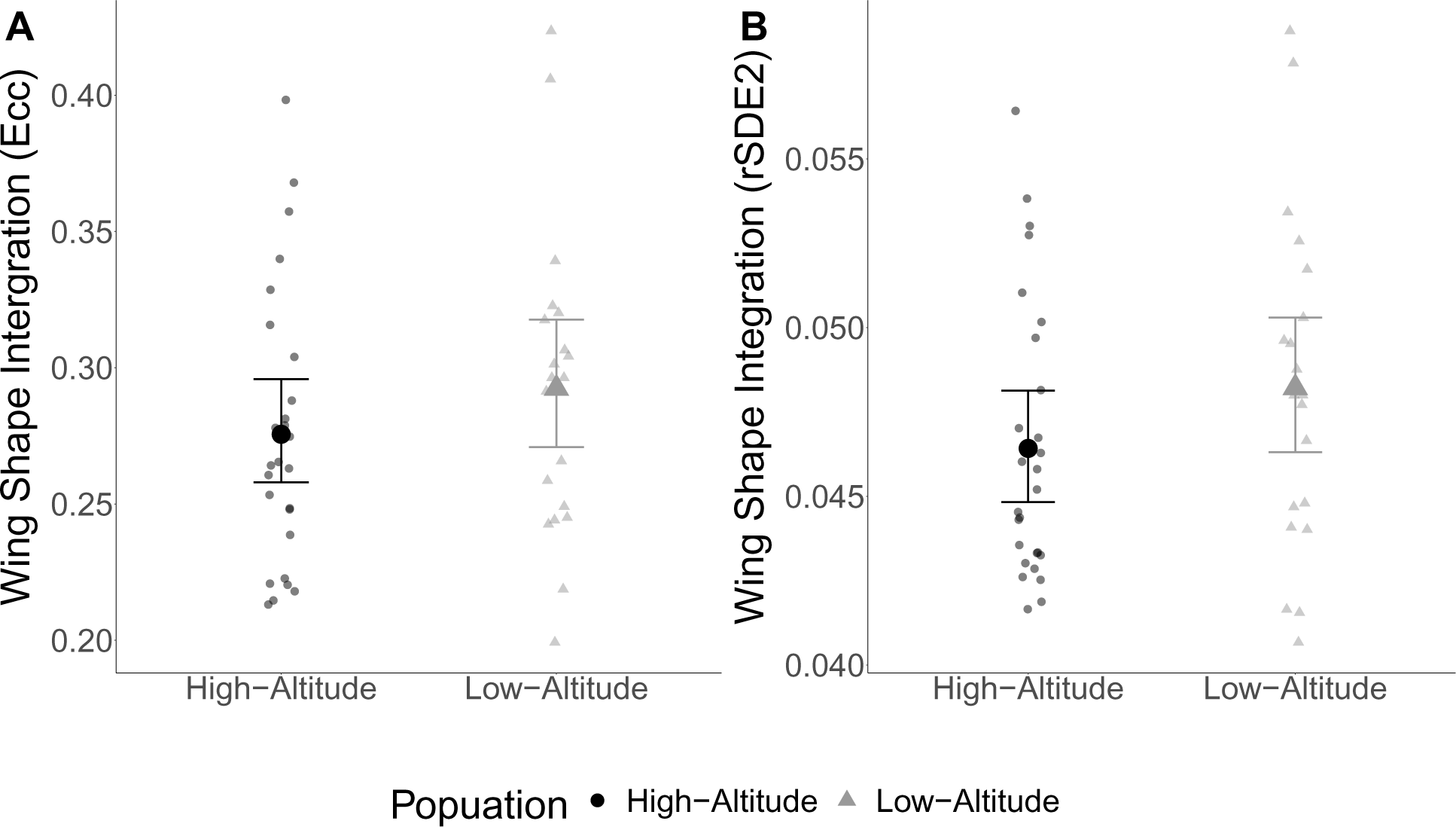
Within-line variation of wing shape measured as wing shape integration using matrix eccentricity and the relative standard deviation (rSDE2) of the eigenvalues of VCV matrix are similar in the high- and low-altitude populations

**Figure S7:**
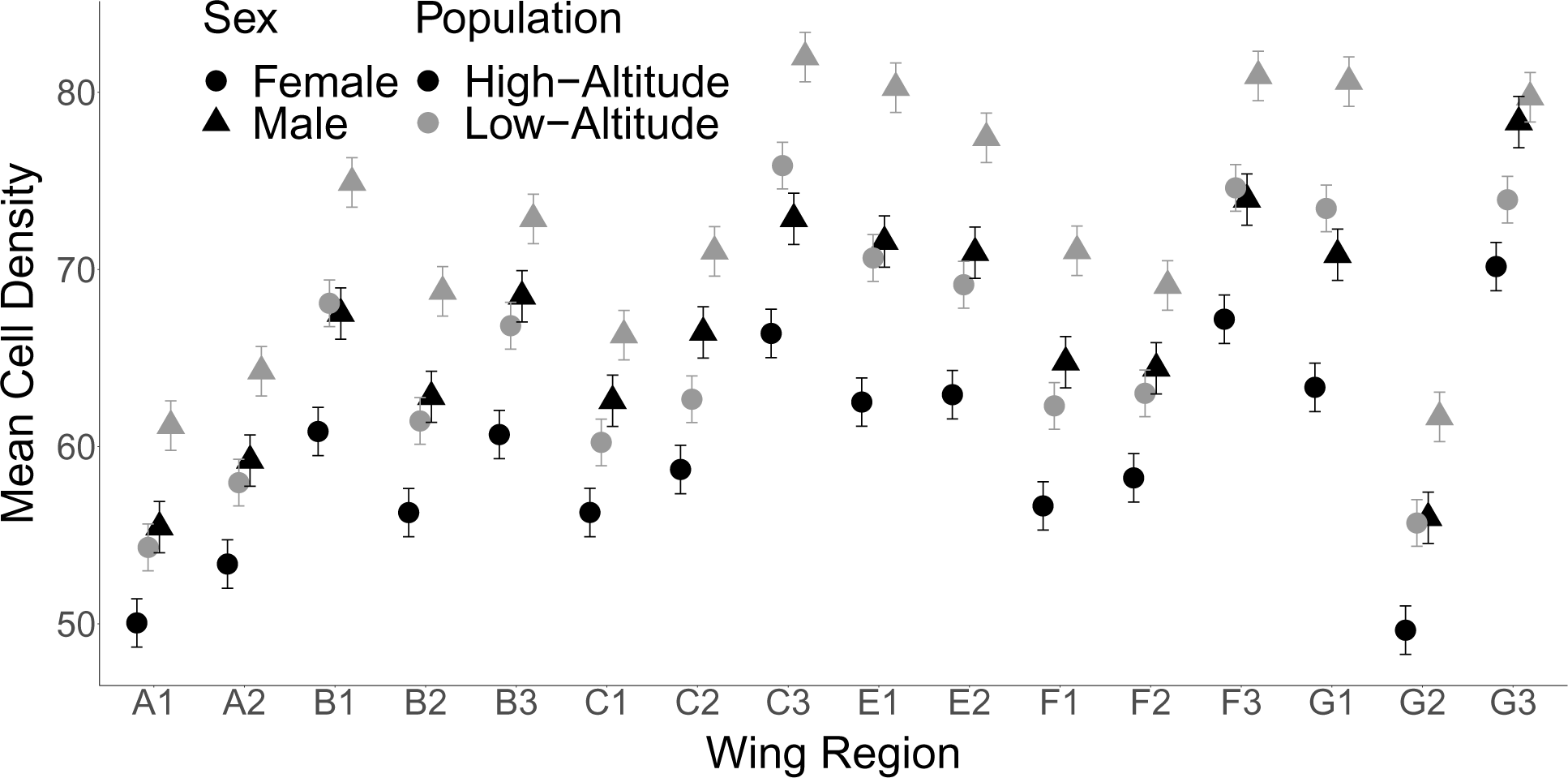
Mean cell density across the 16 measurement wing regions. Wing density varies by wing region, sex and population

**Figure S8:**
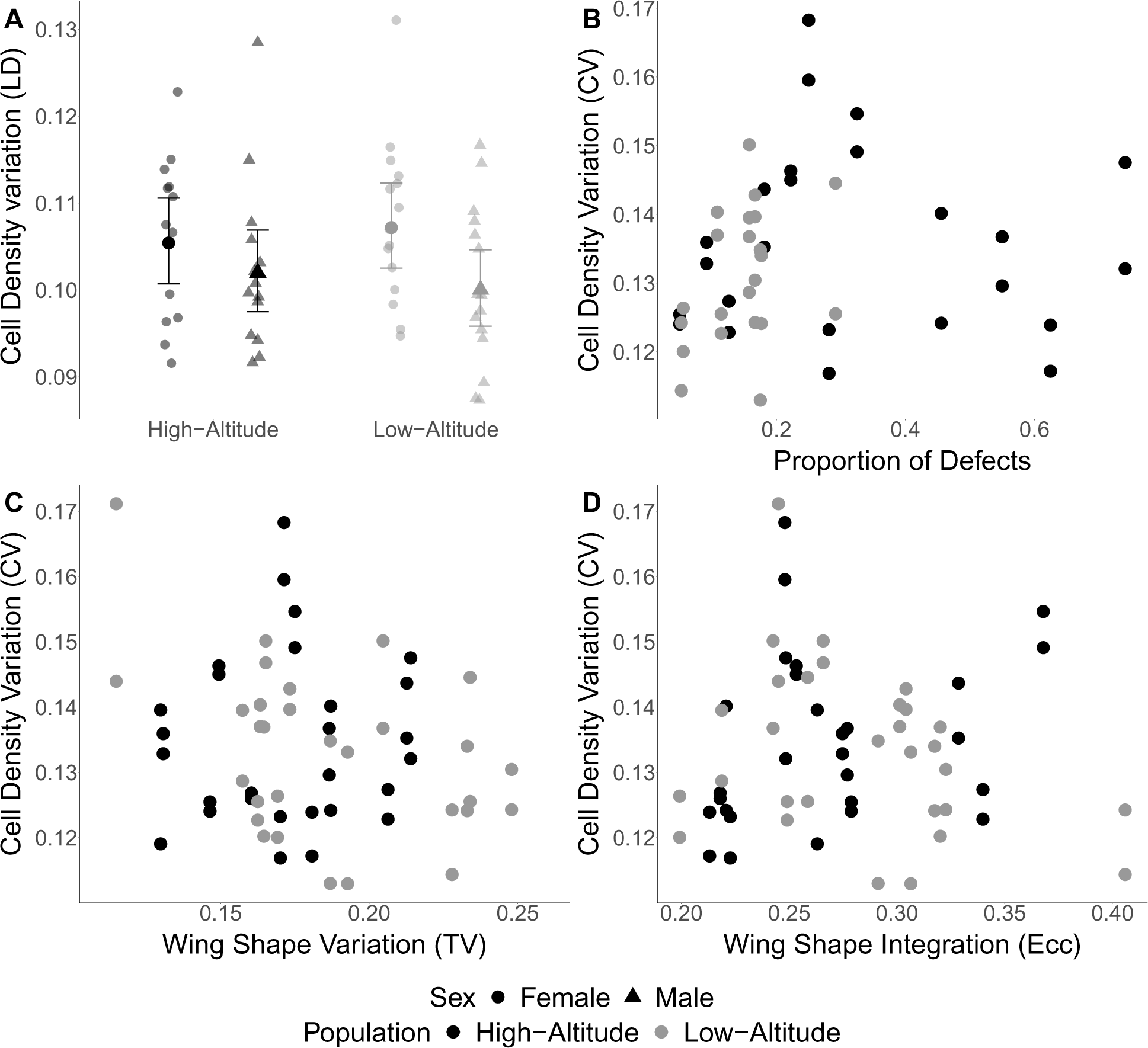
Within-line variation for cell density measured as Levene’s Deviates and association between within-line variation for cell density with within-line wing defects, and within-line variation for wing shape. (A) As seen using CV, within-line variation for cell density using Levene’s deviates is similar between high- and low-altitude populations. (B) There is little association between cell density CV and proportion of defects in either the high-altitude (r = 0.043 95% 0.44 - 0.37) or low-altitude populations (r = 0.36 95% 0.076 - 0.68). Within-line variation (total variance) (C) and integration (eccentricity) (D) for wing shape are not correlated with within-line variation for cell density (TV - Total variance *r* = −0.22 95% CIs 0.45 - 0.04), Ecc - Eccentricity *r* = −0.04 95% CIs 0.29 - 0.22)

**Figure S9:**
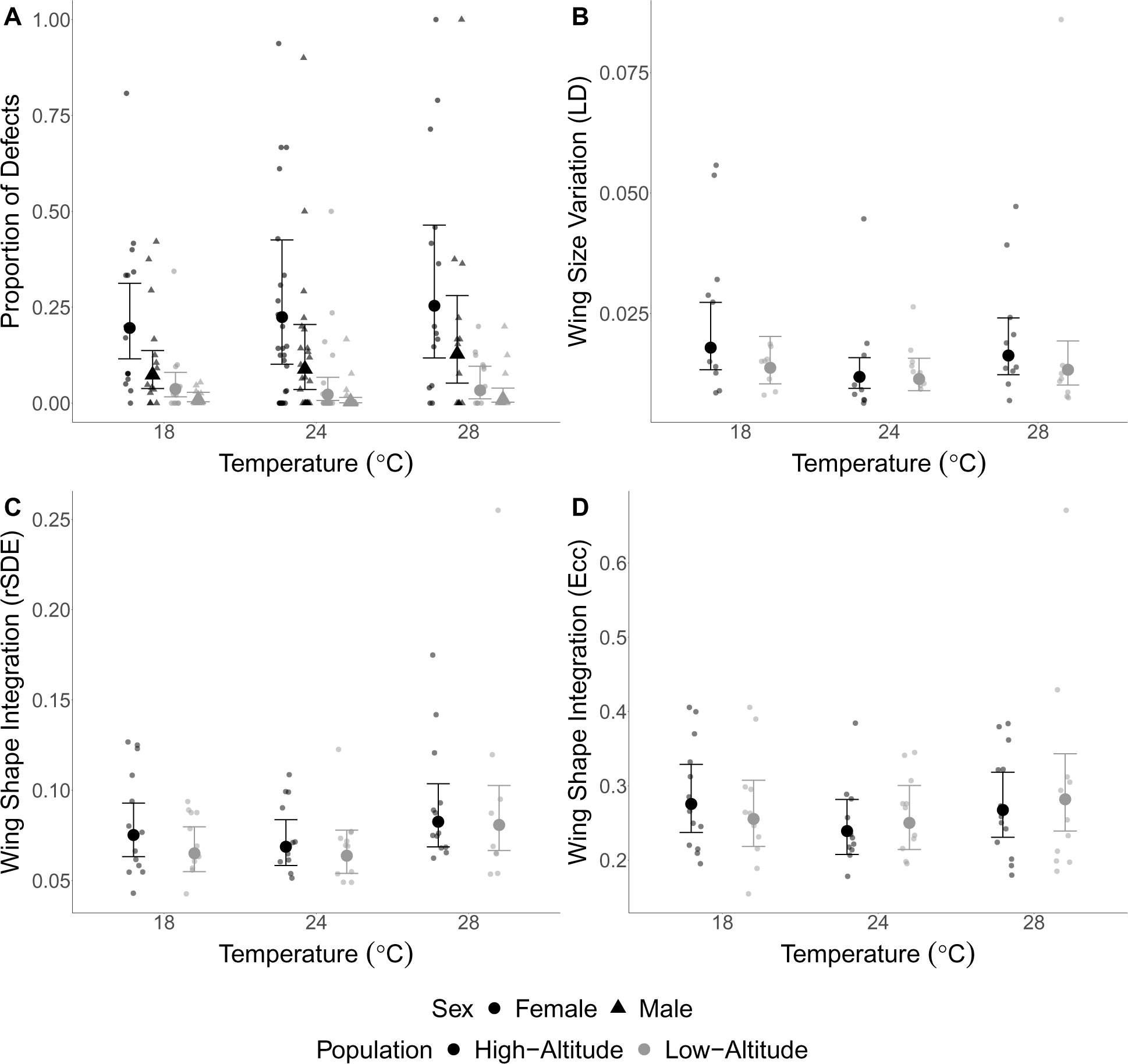
Proportion of wing defects and alternative measures of within-line variation for wing size and shape for the high- and low-altitude population at different developmental temperatures. (A) Proportion of defects are similar across the different developmental temperatures with the high-altitude population having consistently greater proportion of defects than the low-altitude population.(B) Within-line variation for wing size measured as Levene’s Deviates and within-line variation for wing shape measured as (C) rSDE (multiplied by a factor of 10000) and (D) Eccentricity are similar in the high- and low-altitude populations across developmental temperatures.

**Figure S10:**
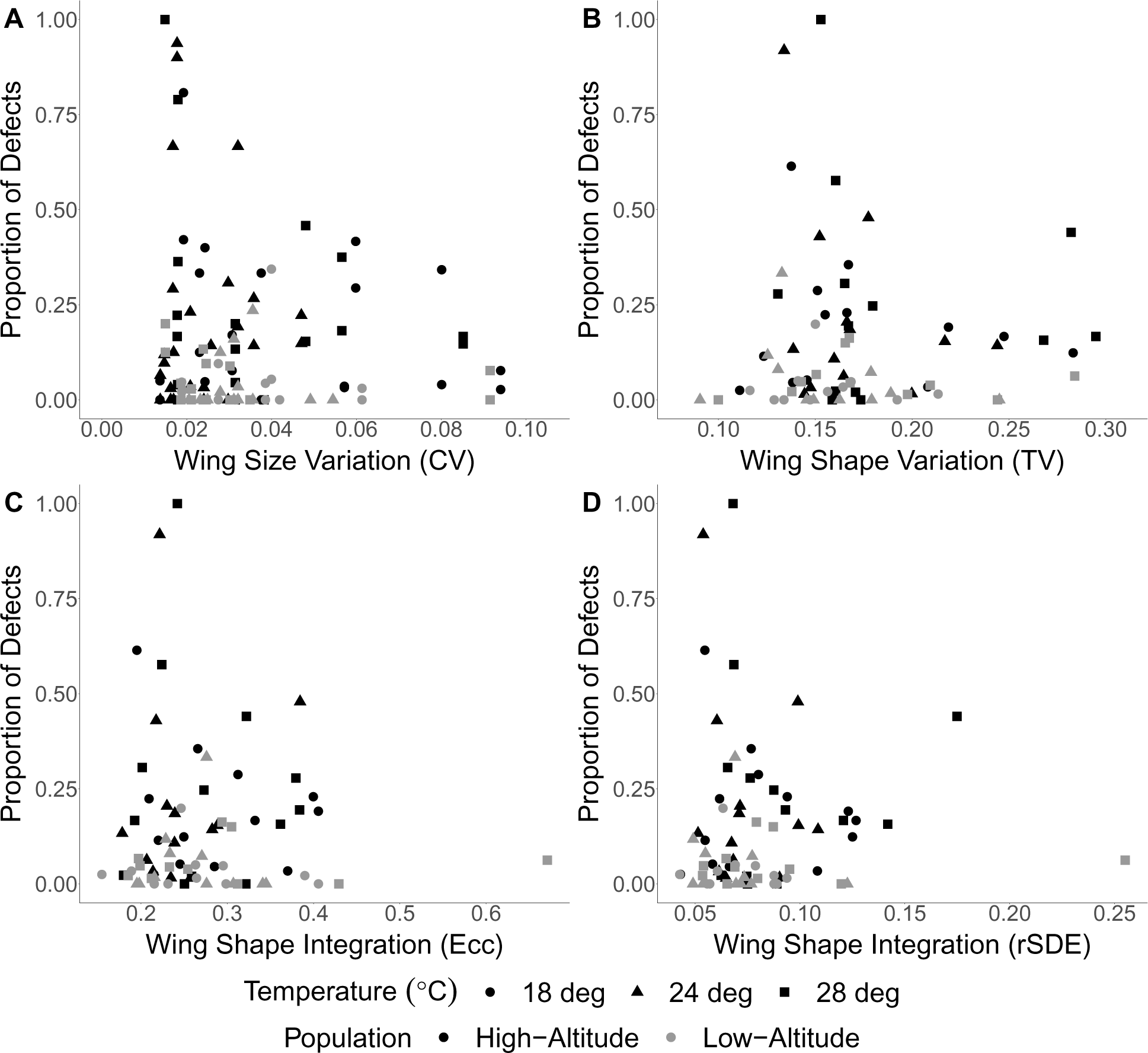
Little evidence for correlation between measures of within-line variation for (A) wing size (CV) and (B-D) wing shape (total variance, eccentricity and rSDE) with proportion of defects for the high- and low-altitude populations at different developmental temperatures

**Figure S11:**
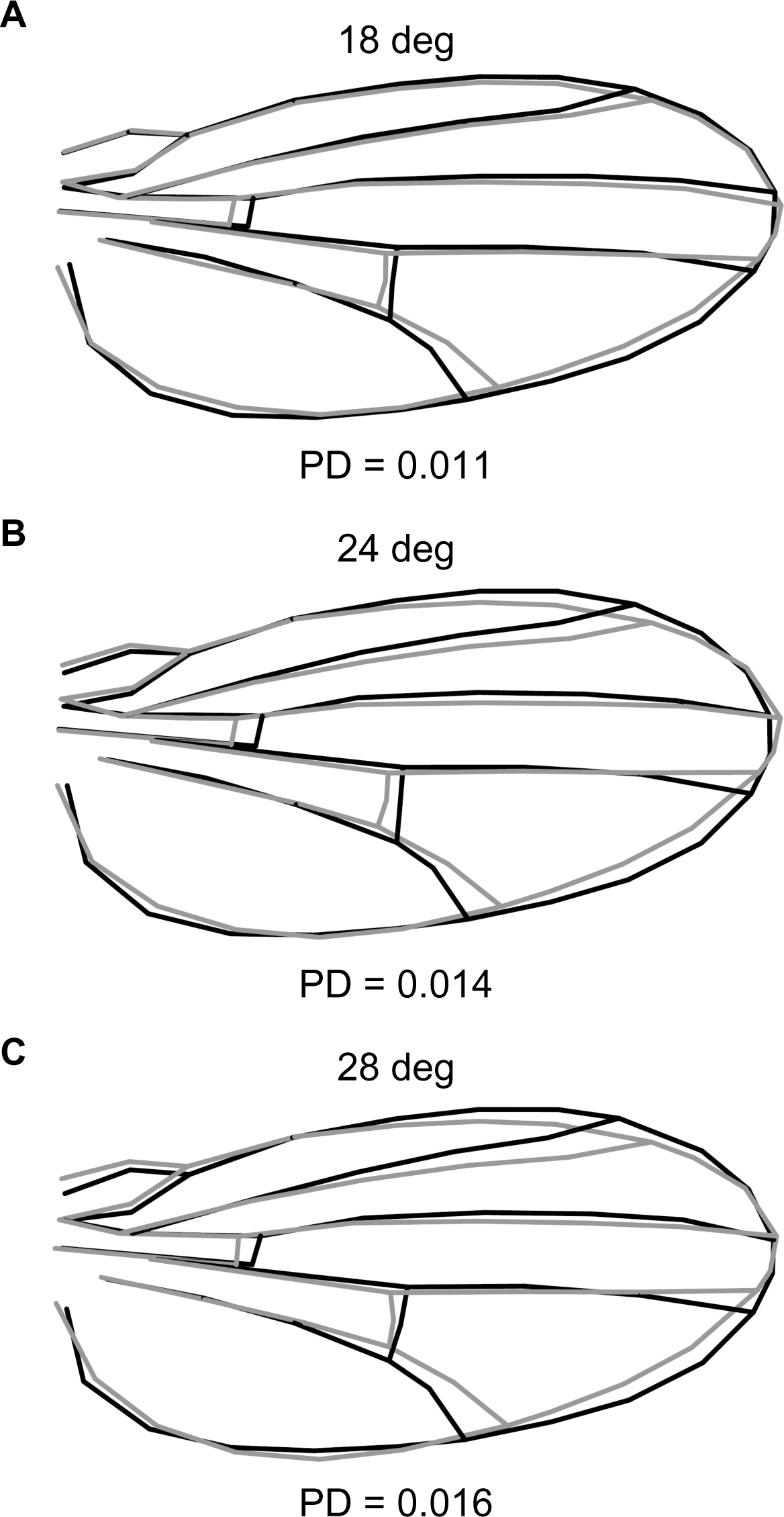
Mean wing shape differences between the high- and low-altitude populations at different temperatures (females). The difference in shape measured as PD (Procrustes distance) is increasing with temperature

**Figure S12:**
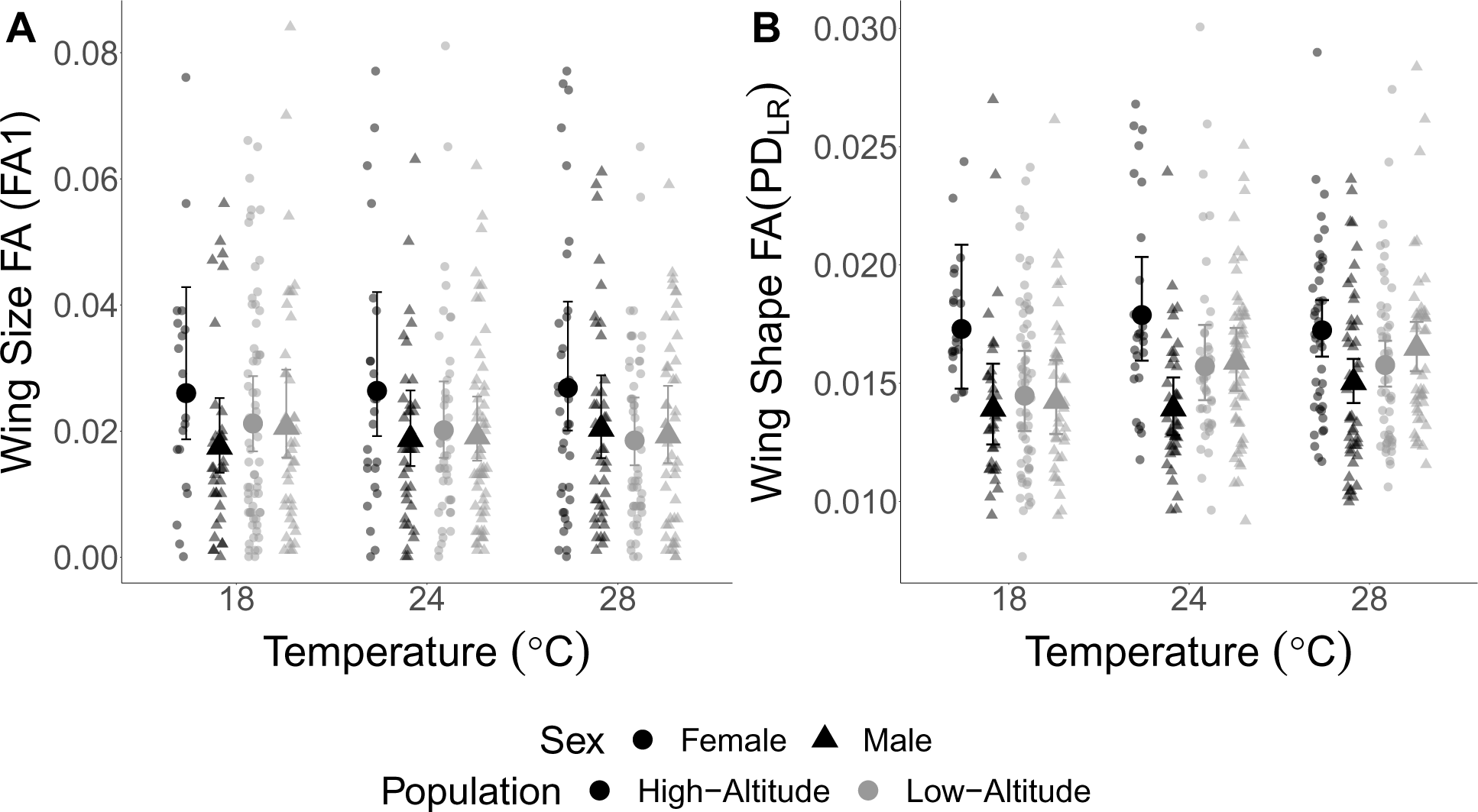
Fluctuating Asymmetry for wing size and shape represented by (A) FA1 and (B) Procrustes distance between the left and right wing (*P D_LR_*). High-altitude females have consistently greater wing size FA and and wing shape FA across the three developmental temperature treatments. Large symbols represent population means, small symbols represent individuals, error bars represent 95% CIs

**Figure S13:**
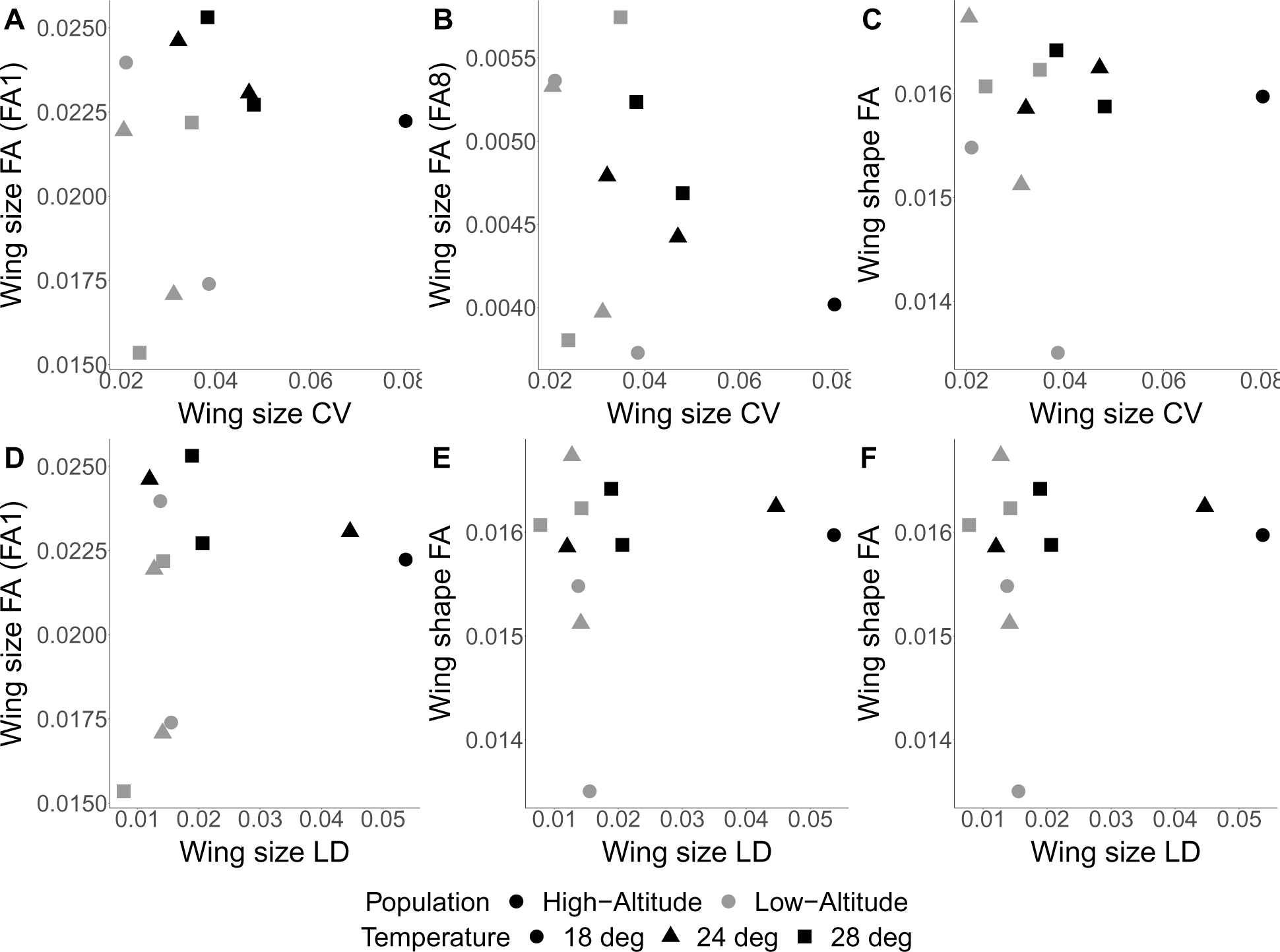
Fluctuating Asymmetry for wing size and shape represented correlated with measures of variability for wing size.(A) FA1 correlated with wing size CV (B) FA8 correlated with wing size CV (C) Procrustes distance between the left and right wing (*P D_LR_*) correlated with wing size CV (D) FA1 correlated with wing size LD (E) FA8 correlated with wing size LD (F)(*P D_LR_*) correlated with wing size LD.

**Figure S14:**
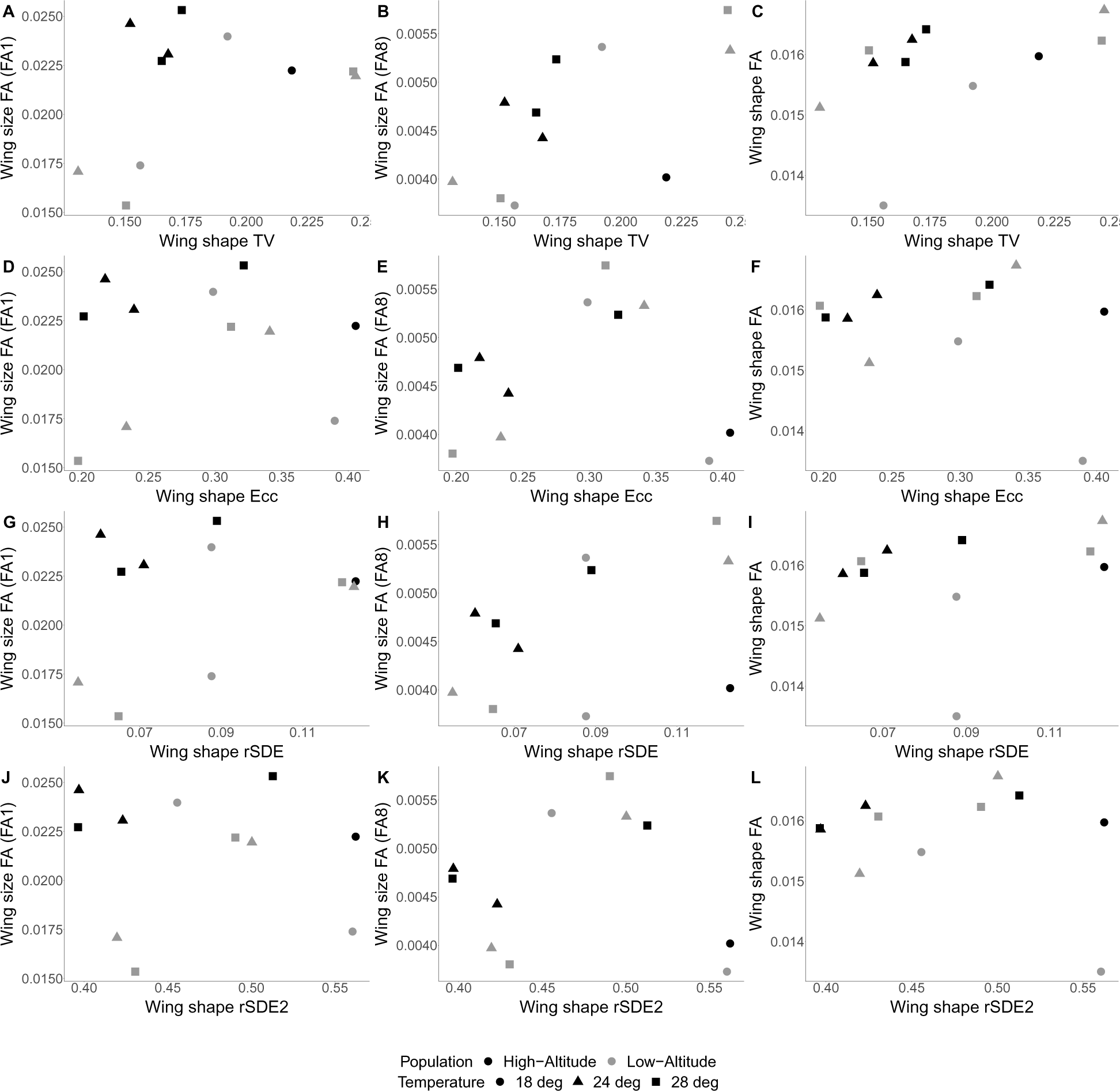
Fluctuating Asymmetry for wing size and shape represented correlated with measures of variability for wing size.(A) FA1 correlated with wing shape total variance (B) FA8 correlated with wing shape total variance (C) Procrustes distance between the left and right wing (*P D_LR_*) correlated with wing shape total variance (D) FA1 correlated with wing shape eccentricity (E) FA8 correlated with wing shape eccentricity (F)(*P D_LR_*) correlated with wing shape eccentricity (G) FA1 correlated with wing shape rSDE (H) FA8 correlated with wing shape rSDE (I)(*P D_LR_*) correlated with wing shape rSDE (J) FA1 correlated with wing shape rSDE2 (K) FA8 correlated with wing shape rSDE2 (L) *P D_LR_*) correlated with wing shape rSDE2.

## Supplemental Tables

**Table S1:** Sample Sizes of fly strains for micro-environmental canalization

| Line | Population | $N_F$ | $N_M$ | Line | Population | $N_F$ | $N_M$ |
| --- | --- | --- | --- | --- | --- | --- | --- |
| ef101 | High-Altitude | 19 | 20 | zi124 | Low-Altitude | 20 | 18 |
| ef112 | High-Altitude | 20 | 20 | zi159 | Low-Altitude | 18 | 15 |
| ef115 | High-Altitude | 19 | 20 | zi160 | Low-Altitude | 17 | 12 |
| ef117 | High-Altitude | 20 | 19 | zi186 | Low-Altitude | 17 | 17 |
| ef119 | High-Altitude | 18 | 14 | zi197 | Low-Altitude | 19 | 19 |
| ef122 | High-Altitude | 18 | 20 | zi216 | Low-Altitude | 16 | 17 |
| ef131 | High-Altitude | 20 | 20 | zi217 | Low-Altitude | 19 | 20 |
| ef135 | High-Altitude | 19 | 20 | zi251 | Low-Altitude | 19 | 14 |
| ef136 | High-Altitude | 16 | 20 | zi254 | Low-Altitude | 12 | 17 |
| ef15 | High-Altitude | 15 | 20 | zi311 | Low-Altitude | 17 | 19 |
| ef16 | High-Altitude | 20 | 19 | zi322 | Low-Altitude | 23 | 14 |
| ef19 | High-Altitude | 20 | 19 | zi337 | Low-Altitude | 19 | 20 |
| ef1 | High-Altitude | 20 | 17 | zi357 | Low-Altitude | 20 | 20 |
| ef25 | High-Altitude | 20 | 20 | zi360 | Low-Altitude | 18 | 14 |
| ef31 | High-Altitude | 20 | 19 | zi366 | Low-Altitude | 18 | 20 |
| ef39 | High-Altitude | 20 | 20 | zi383 | Low-Altitude | 19 | 19 |
| ef3 | High-Altitude | 18 | 14 | zi403 | Low-Altitude | 17 | 16 |
| ef54 | High-Altitude | 20 | 19 | zi455 | Low-Altitude | 20 | 19 |
| ef59 | High-Altitude | 19 | 13 | zi461 | Low-Altitude | 14 | 18 |
| ef65 | High-Altitude | 19 | 19 | zi507 | Low-Altitude | 20 | 18 |
| ef67 | High-Altitude | 11 | 17 |  |  |  |  |
| ef73 | High-Altitude | 18 | 16 |  |  |  |  |
| ef75 | High-Altitude | 8 | 18 |  |  |  |  |
| ef7 | High-Altitude | 14 | 19 |  |  |  |  |
| ef86 | High-Altitude | 19 | 20 |  |  |  |  |
| ef98 | High-Altitude | 14 | 20 |  |  |  |  |
| ef9 | High-Altitude | 17 | 20 |  |  |  |  |

**Table S2:** Sample Sizes of fly strains for Temperature Plasticity and Macro-environmental canalization

| Line | Population | $N_{18^{\circ}CF}$ | $N_{24^{\circ}CF}$ | $N_{28^{\circ}CF}$ | $N_{18^{\circ}CM}$ | $N_{24^{\circ}CM}$ | $N_{28^{\circ}CM}$ |
| --- | --- | --- | --- | --- | --- | --- | --- |
| ef112 | High-Altitude | 28 | 10 | 22 | 18 | 23 | 9 |
| ef117 | High-Altitude | 7 | 22 | 8 | 9 | 5 | 8 |
| ef119 | High-Altitude | 20 | 24 | 11 | 16 | 9 | 8 |
| ef122 | High-Altitude | 8 | 14 | 6 | 11 | 19 | 18 |
| ef131 | High-Altitude | 26 | 27 | 41 | 35 | 39 | 22 |
| ef136 | High-Altitude | 12 | 20 | 16 | 14 | 20 | 10 |
| ef15 | High-Altitude | 13 | 19 | 21 | 19 | 24 | 14 |
| ef19 | High-Altitude | 7 | 12 | 9 | 6 | 13 | 10 |
| ef39 | High-Altitude | 7 | 13 | 17 | 12 | 21 | 23 |
| ef65 | High-Altitude | 13 | 20 | 4 | 8 | 14 | 6 |
| ef73 | High-Altitude | 36 | 21 | 12 | 23 | 26 | 8 |
| ef16 | High-Altitude | 42 | 27 | 43 | 36 | 26 | 30 |
| ef98 | High-Altitude | 25 | 16 | 13 | 19 | 20 | 8 |
| zi124 | Low-Altitude | 47 | 20 | 31 | 39 | 21 | 25 |
| zi159 | Low-Altitude | 41 | 72 | 42 | 39 | 60 | 31 |
| zi186 | Low-Altitude | 26 | 8 | 21 | 37 | 8 | 23 |
| zi216 | Low-Altitude | 26 | 47 | 10 | 29 | 56 | 4 |
| zi251 | Low-Altitude | 20 | 32 | 17 | 23 | 47 | 28 |
| zi254 | Low-Altitude | 40 | 40 | 40 | 38 | 49 | 36 |
| zi311 | Low-Altitude | 22 | 22 | 15 | 13 | 16 | 12 |
| zi360 | Low-Altitude | 9 | 17 | 5 | 20 | 9 | 4 |
| zi366 | Low-Altitude | 4 | 12 | 8 | 14 | 6 | 2 |
| zi367 | Low-Altitude | 8 | 35 | 23 | 8 | 23 | 10 |
| zi455 | Low-Altitude | 32 | 32 | 15 | 35 | 37 | 16 |

**Table S3:**
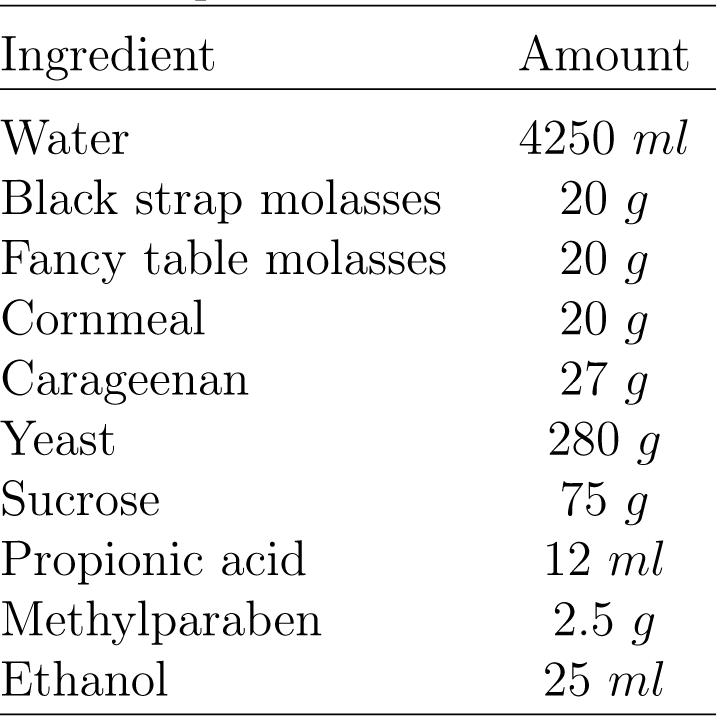
Recipe for 1.5:1 Protein:Sugar Food used in Temperature Plasticity/Macro-environmental canalization and FA experiments

| Ingredient | Amount |
| --- | --- |
| Water | 4250 *ml* |
| Black strap molasses | 20 *g* |
| Fancy table molasses | 20 *g* |
| Cornmeal | 20 *g* |
| Carageenan | 27 *g* |
| Yeast | 280 *g* |
| Sucrose | 75 *g* |
| Propionic acid | 12 *ml* |
| Methylparaben | 2.5 *g* |
| Ethanol | 25 *ml* |

**Table S4:** Linear mixed estimates for the contributions of sex, population and their interaction on wing size

| Effect | Chisq | Df | Pr(>Chisq) |
| --- | --- | --- | --- |
| Sex | $4.6 \times 10^3$ | 1 | $< 2.2 \times 10^{-16}$ |
| Pop | $9.5 \times 10^2$ | 1 | $< 2.2 \times 10^{-16}$ |
| Sex:Pop | $1.0 \times 10^2$ | 1 | $< 2.2 \times 10^{-16}$ |

**Table S5:** Results from Multivariate Procrustes ANOVA testing the effects of wing size, population, sex and all interactions on wing shape

| Effect | Df | SS | MS | Rsqr | F | Z | p (>F) |
| --- | --- | --- | --- | --- | --- | --- | --- |
| Size | 1 | $5.7 \times 10^{-2}$ | $5.7 \times 10^{-2}$ | $5.8 \times 10^{-2}$ | $2.7 \times 10^2$ | 11 | $5.0 \times 10^{-4}$ |
| Pop | 1 | $1.1 \times 10^{-1}$ | $1.1 \times 10^{-1}$ | $1.1 \times 10^{-1}$ | 11 | 4.7 | $5.0 \times 10^{-4}$ |
| Sex | 1 | $1.0 \times 10^{-2}$ | $1.0 \times 10^{-2}$ | $1.0 \times 10^{-2}$ | 47 | 9 | $5.0 \times 10^{-4}$ |
| Size:Pop | 1 | $3.3 \times 10^{-3}$ | $3.3 \times 10^{-3}$ | $3.3 \times 10^{-3}$ | 16 | 6.4 | $5.0 \times 10^{-4}$ |
| Size:Sex | 1 | $1.3 \times 10^{-3}$ | $1.3 \times 10^{-3}$ | $1.3 \times 10^{-3}$ | 6.0 | 4.5 | $5.0 \times 10^{-4}$ |
| Pop:Sex | 1 | $5.5 \times 10^{-4}$ | $5.5 \times 10^{-4}$ | $5.6 \times 10^{-4}$ | 2.6 | 2.7 | $5.0 \times 10^{-4}$ |
| Pop:Line | 45 | $4.5 \times 10^{-1}$ | $1.0 \times 10^{-2}$ | $4.6 \times 10^{-1}$ | 47 | 35 | $5.0 \times 10^{-4}$ |
| Size:Pop:Sex | 1 | $4.0 \times 10^{-4}$ | $3.9 \times 10^{-4}$ | $4.0 \times 10^{-4}$ | 1.8 | 4.0 | $5.0 \times 10^{-4}$ |
| Residuals | $1.6 \times 10^3$ | $3.5 \times 10^{-1}$ | $2.1 \times 10^{-4}$ | $3.5 \times 10^{-1}$ | | | |
| Total | $1.7 \times 10^3$ | $9.8 \times 10^{-1}$ | | | | | |

**Table S6:** Results from generalized mixed effects model testing the effects of population and sex and their interaction on within-line among-individual wing size CV

| Effect | Chisq | Df | Pr(>Chisq) |
| --- | --- | --- | --- |
| Pop | 0.13 | 1 | 0.72 |
| Sex | $1.0 \times 10^{-3}$ | 1 | 0.98 |
| Pop:Sex | 0.41 | 1 | 0.52 |

**Table S7:** Results from generalized mixed effects model testing the effects of sex and population and their interaction on within-line among-individual wing size Levene’s deviates

| Effect | Chisq | Df | Pr(>Chisq) |
| --- | --- | --- | --- |
| Sex | 0.84 | 1 | 0.36 |
| Pop | 1.5 | 1 | 0.22 |
| Sex:Pop | 0.20 | 1 | 0.66 |

**Table S8:**
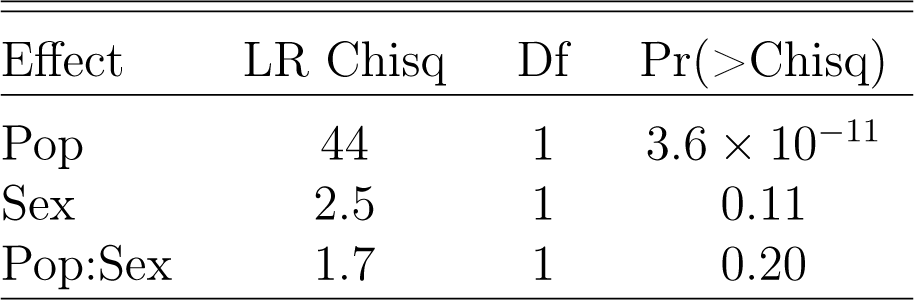
Results from generalized mixed effects model testing the effects of population and sex and their interaction on within-line proportion of wing defects

**Table S9:** Results from generalized mixed effects model testing the effects of population on within-line total variance for wing shape

| Effect | LR Chisq | Df | Pr(>Chisq) |
| --- | --- | --- | --- |
| Pop | $5.6 \times 10^3$ | 1 | 0.94 |

**Table S10:** Results from generalized mixed effects model testing the effects of population on within-line eccentricity for wing shape

| Effect | LR Chisq | Df | Pr(>Chisq) |
| --- | --- | --- | --- |
| Pop | 1.3 | 1 | 0.25 |

**Table S11:** Results from linear mixed effects model testing the effects of wing region, sex, population and their interactions on cell density across 16 different regions of the wing

| Effect | Chisq | Df | Pr(>Chisq) |
| --- | --- | --- | --- |
| WingReg | $1.9 \times 10^4$ | 15 | $< 2.2 \times 10^{-16}$ |
| Sex | $5.6 \times 10^2$ | 1 | $< 2.2 \times 10^{-16}$ |
| Pop | 31 | 1 | $2.0 \times 10^{-8}$ |
| WingReg:Sex | $1.1 \times 10^2$ | 15 | $< 2.2 \times 10^{-16}$ |
| WingReg:Pop | $4.0 \times 10^2$ | 15 | $< 2.2 \times 10^{-16}$ |
| Sex:Pop | 0.31 | 1 | 0.58 |
| WingReg:Sex:Pop | 20 | 15 | 0.16 |

**Table S12:** Results from generalized mixed effects model testing the effects of population and sex and their interaction on within-line among-individual cell density CV

| Effect | Chisq | Df | Pr(>Chisq) |
| --- | --- | --- | --- |
| Pop | $4.2 \times 10^{-2}$ | 1 | 0.84 |
| Sex | 10 | 1 | $1.0 \times 10^{-3}$ |
| Pop:Sex | 0.27 | 1 | 0.60 |

**Table S13:** Results from generalized mixed effects model testing the effects of population and sex and their interaction on within-line among-individual cell density Levene’s deviates

| Effect | Chisq | Df | Pr(>Chisq) |
| --- | --- | --- | --- |
| Sex | 14 | 1 | $2.0 \times 10^{-4}$ |
| Pop | $3.0 \times 10^{-3}$ | 1 | 0.95 |
| Sex:Pop | 0.63 | 1 | 0.43 |

**Table S14:**
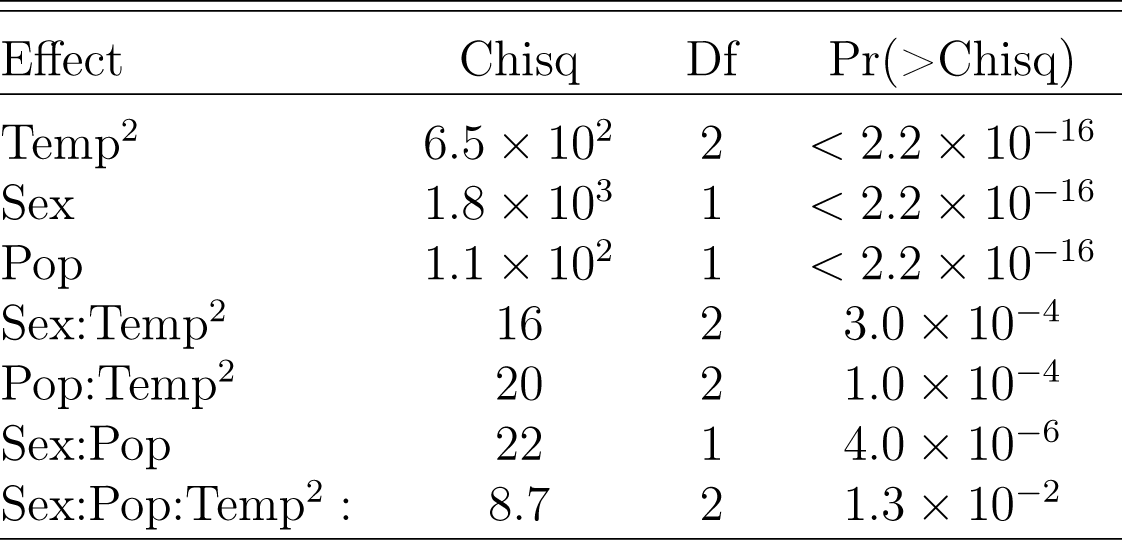
Results from linear mixed effects model testing the effects of temperature, sex, population and their interactions on wing size

**Table S15:** Results from Multivariate Procrustes ANOVA testing the effects of wing size, sex, temperature, population and all interactions on wing shape

| Effect | Df | SS | MS | Rsqr | F | Z | p (>F) |
| --- | --- | --- | --- | --- | --- | --- | --- |
| Size | 1 | $7.6 \times 10^{-2}$ | $7.6 \times 10^{-2}$ | $3.6 \times 10^{-2}$ | $3.4 \times 10^2$ | 12 | $5.0 \times 10^{-4}$ |
| Sex | 1 | 0.14 | 0.14 | $6.7 \times 10^{-2}$ | $6.3 \times 10^2$ | 13 | $5.0 \times 10^{-4}$ |
| Temp | 1 | 0.13 | 0.13 | $6.1 \times 10^{-2}$ | $5.8 \times 10^2$ | 13 | $5.0 \times 10^{-4}$ |
| Temp<sup>2</sup> | 1 | $2.0 \times 10^{-2}$ | $2.0 \times 10^{-2}$ | $9.4 \times 10^{-3}$ | 89 | 9.6 | $5.0 \times 10^{-4}$ |
| Pop | 1 | $5.90 \times 10^{-2}$ | $5.9 \times 10^{-2}$ | $2.8 \times 10^{-2}$ | 1.7 | 1.1 | 0.14 |
| Size:Sex | 1 | $2.6 \times 10^{-3}$ | $2.6 \times 10^{-3}$ | $1.5 \times 10^{-3}$ | 12 | 5.8 | $5.0 \times 10^{-4}$ |
| Size:Temp | 1 | $3.8 \times 10^{-3}$ | $3.8 \times 10^{-3}$ | $1.8 \times 10^{-3}$ | 17 | 6.4 | $5.0 \times 10^{-4}$ |
| Size:Temp<sup>2</sup> | 1 | $1.4 \times 10^{-3}$ | $1.4 \times 10^{-3}$ | $6.6 \times 10^{-4}$ | 6.2 | 4.4 | $5.0 \times 10^{-4}$ |
| Size:Pop | 1 | $2.0 \times 10^{-2}$ | $2.0 \times 10^{-2}$ | $9.4 \times 10^{-3}$ | 89 | 10 | $5.0 \times 10^{-4}$ |
| Sex:Temp | 1 | $3.9 \times 10^{-3}$ | $3.9 \times 10^{-3}$ | $1.9 \times 10^{-3}$ | 18 | 6.5 | $5.0 \times 10^{-4}$ |
| Sex:Temp<sup>2</sup> | 1 | $1.5 \times 10^{-3}$ | $1.5 \times 10^{-3}$ | $7.3 \times 10^{-4}$ | 6.9 | 4.7 | $5.0 \times 10^{-4}$ |
| Sex:Pop | 1 | $6.9 \times 10^{-3}$ | $6.9 \times 10^{-3}$ | $3.3 \times 10^{-3}$ | 31 | 7.5 | $5.0 \times 10^{-4}$ |
| Pop:Temp | 1 | $1.3 \times 10^{-2}$ | $1.3 \times 10^{-2}$ | $6.2 \times 10^{-3}$ | 59 | 9.0 | $5.0 \times 10^{-4}$ |
| Pop:Temp<sup>2</sup> | 1 | $4.4 \times 10^{-3}$ | $4.4 \times 10^{-3}$ | $2.1 \times 10^{-3}$ | 20 | 6.8 | $5.0 \times 10^{-4}$ |
| Pop:Line | 25 | 0.89 | $3.5 \times 10^{-2}$ | 0.42 | $1.6 \times 10^2$ | 42 | $5.0 \times 10^{-4}$ |
| Size:Sex:Temp | 1 | $5.0 \times 10^{-4}$ | $5.0 \times 10^{-4}$ | $2.4 \times 10^{-4}$ | 2.0 | 4.4 | $5.0 \times 10^{-4}$ |
| Size:Sex:Temp<sup>2</sup> | 1 | $4.5 \times 10^{-4}$ | $4.5 \times 10^{-4}$ | $2.1 \times 10^{-4}$ | 2.2 | 4.7 | $5.0 \times 10^{-4}$ |
| Size:Sex:Pop | 1 | $1.4 \times 10^{-3}$ | $1.4 \times 10^{-3}$ | $6.8 \times 10^{-4}$ | 6.4 | 7.2 | $5.0 \times 10^{-4}$ |
| Size:Pop:Temp | 1 | $6.0 \times 10^{-4}$ | $6.0 \times 10^{-4}$ | $2.9 \times 10^{-4}$ | 2.7 | 5.2 | $5.0 \times 10^{-4}$ |
| Size:Pop:Temp<sup>2</sup> | 1 | $1.7 \times 10^{-3}$ | $1.7 \times 10^{-3}$ | $8.1 \times 10^{-4}$ | 7.6 | 7.6 | $5.0 \times 10^{-4}$ |
| Sex:Pop:Temp | 1 | $9.8 \times 10^{-4}$ | $9.8 \times 10^{-4}$ | $4.7 \times 10^{-4}$ | 4.4 | 6.4 | $5.0 \times 10^{-4}$ |
| Sex:Pop:Temp<sup>2</sup> | 1 | $3.2 \times 10^{-4}$ | $3.2 \times 10^{-4}$ | $1.5 \times 10^{-4}$ | 1.4 | 3.7 | $5.0 \times 10^{-4}$ |
| Residuals | $3.3 \times 10^3$ | 0.73 | $2.2 \times 10^{-4}$ | 0.35 | | | |
| Total | $3.3 \times 10^3$ | 2.1 | | | | | |

**Table S16:** Results from generalized mixed effects model testing the effects population, temperature and their interaction on within-line among-individual CV for wing size

|  | Chisq | Df | Pr(>Chisq) |
| --- | --- | --- | --- |
| Pop | $5.0 \times 10^{-4}$ | 1 | 0.99 |
| Temp | 8.1 | 2 | $1.8 \times 10^{-2}$ |
| Pop:Temp | 5.2 | 2 | $7.5 \times 10^{-2}$ |

**Table S17:**
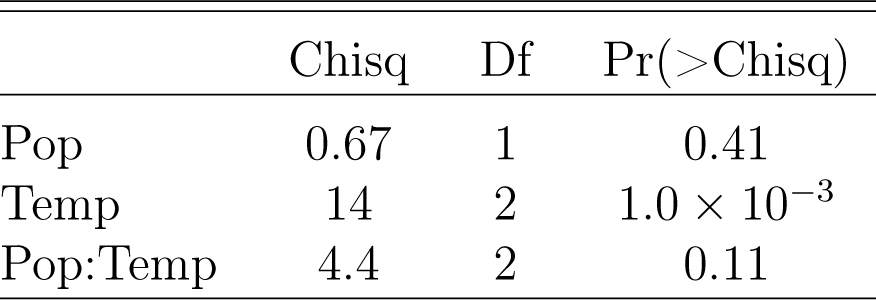
Results from generalized mixed effects model testing the effects of population, temperature and thier interaction on within-line among-individual Levene’s deviates for wing size

**Table S18:** Results from generalized mixed effects model testing the effects of population, temperature and their interaction on within-line among-individual total variance for wing shape

| Effect | Chisq | Df | Pr(>Chisq) |
| --- | --- | --- | --- |
| Pop | 0.44 | 1 | 0.51 |
| Temp | 10 | 2 | $6.0 \times 10^{-3}$ |
| Pop:Temp | 0.85 | 2 | 0.65 |

**Table S19:** Results from generalized mixed effects model testing the effects of population, temperature and their interaction on within-line among-individual eccentricity for wing shape

| Effect | Chisq | Df | Pr(>Chisq) |
| --- | --- | --- | --- |
| Pop | $4.0 \times 10^{-3}$ | 1 | 0.95 |
| Temp | 3.7 | 2 | 0.15 |
| Pop:Temp | 0.91 | 2 | 0.63 |

**Table S20:**
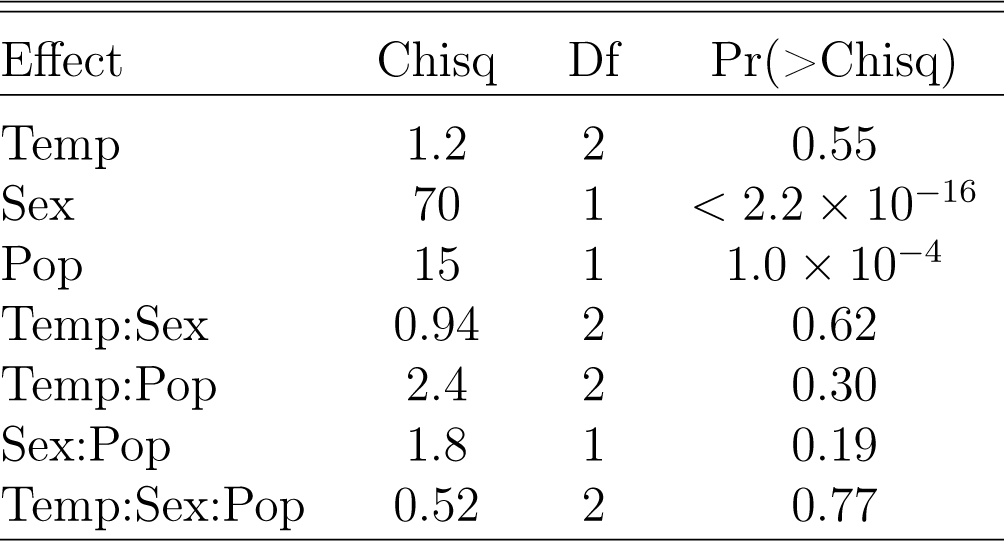
Results from generalized mixed effects model testing the effects of temperature, populations sex and all interactions on within-line proportion of wing defects

| Effect | Chisq | Df | Pr(>Chisq) |
| --- | --- | --- | --- |
| Temp | 1.2 | 2 | 0.55 |
| Sex | 70 | 1 | $< 2.2 \times 10^{-16}$ |
| Pop | 15 | 1 | $1.0 \times 10^{-4}$ |
| Temp:Sex | 0.94 | 2 | 0.62 |
| Temp:Pop | 2.4 | 2 | 0.30 |
| Sex:Pop | 1.8 | 1 | 0.19 |
| Temp:Sex:Pop | 0.52 | 2 | 0.77 |

**Table S21:** Results from generalized mixed effects model testing the effects of sex, population, temperature and all interactions on FA1

| Effect | Chisq | Df | Pr(>Chisq) |
| --- | --- | --- | --- |
| Sex | 3.2 | 1 | $7.2 \times 10^{-2}$ |
| Pop | 1.1 | 1 | 0.28 |
| Temp | 0.11 | 2 | 0.94 |
| Sex:Pop | 3.9 | 1 | $4.7 \times 10^{-2}$ |
| Sex:Temp | 0.34 | 2 | 0.84 |
| Pop:Temp | 1.0 | 2 | 0.60 |
| Sex:Pop:Temp | $9.4 \times 10^{-2}$ | 2 | 0.95 |

**Table S22:** Results from generalized mixed effects model testing the effects of sex, population, temperature and all interactions on FA8

| Effect | Chisq | Df | Pr(>Chisq) |
| --- | --- | --- | --- |
| Sex | $9.0 \times 10^{-3}$ | 1 | 0.92 |
| Pop | $1.0 \times 10^{-2}$ | 1 | 0.92 |
| Temp | 1.0 | 2 | 0.60 |
| Sex:Pop | 10 | 1 | $1.0 \times 10^{-3}$ |
| Sex:Temp | 0.80 | 2 | 0.67 |
| Pop:Temp | 2.4 | 2 | 0.29 |
| Sex:Pop:Temp | 0.84 | 2 | 0.66 |

**Table S23:**
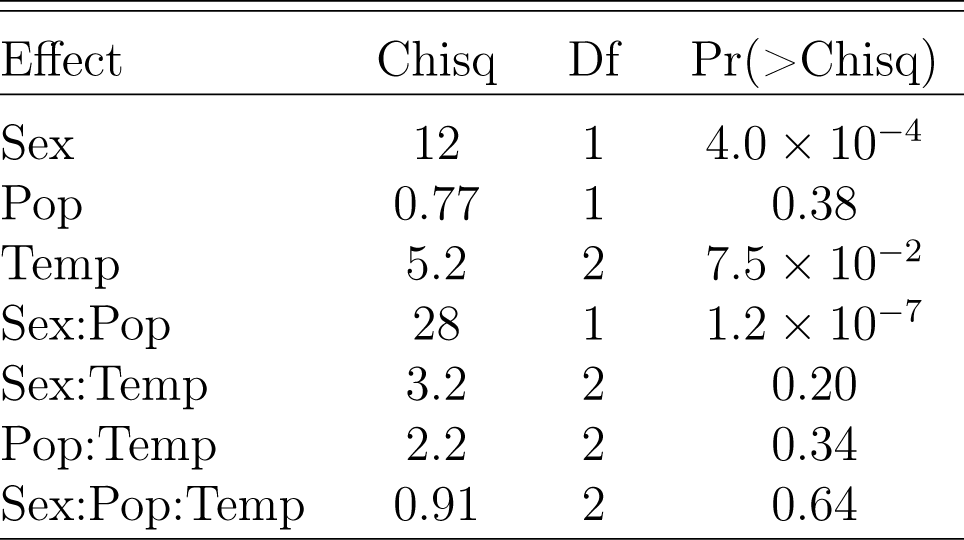
Results from generalized mixed effects model testing the effects of sex, population, temperature and all interactions on *P D_LR_*

**Table S24:** Results from generalized model testing the effects of wing size, sex, population, temperature and all interactions on wing shape FA after correcting for DA

|  | LR Chisq | Df | Pr(>Chisq) |
| --- | --- | --- | --- |
| Pop | 1.4 | 1 | 0.24 |
| Sex | 4.1 | 1 | 0.043 |
| Temp | 0.90 | 2 | 0.64 |
| Size | 3.2 | 1 | 0.075 |
| Pop:Sex | 1.3 | 1 | 0.25 |
| Pop:Temp | 2.0 | 2 | 0.37 |
| Pop:Size | 1.3 | 1 | 0.26 |
| Sex:Temp | 1.6 | 2 | 0.44 |
| Sex:Size | 0.070 | 1 | 0.79 |
| Temp:Size | 1.0 | 2 | 0.60 |
| Pop:Sex:Temp | 0.15 | 2 | 0.93 |
| Pop:Sex:Size | 2.7 | 1 | 0.10 |
| Pop:Temp:Size | 8.8 | 2 | 0.012 |
| Sex:Temp:Size | 2.1 | 2 | 0.36 |

**Table S25:** Results from wing size ANOVA on repeated measurements using side, individual and their interaction as effects in order to estimate measurement error

| Effect | Sum Sq | Df | F value | Pr(>F) |
| --- | --- | --- | --- | --- |
| Side | $4.0 \times 10^{-4}$ | 1 | 14 | $3.0 \times 10^{-4}$ |
| Inds | $1.0 \times 10^2$ | 76 | $4.2 \times 10^4$ | $< 2.2 \times 10^{-16}$ |
| Side:Inds | 0.31 | 76 | $1.3 \times 10^2$ | $< 2.2 \times 10^{-16}$ |
| Residuals | $5.0 \times 10^{-3}$ | 154 | | |

**Table S26:** Results from wing shape multivariate procrustes ANOVA on repeated measurements using side, individual and their interaction as effects in order to estimate Wing shape Measurement Error ANOVA

| Effect | Df | SS | MS | Rsq | F | Z | Pr(>F) |
| --- | --- | --- | --- | --- | --- | --- | --- |
| Side | 1 | $8.7 \times 10^{-4}$ | $8.7 \times 10^{-4}$ | $5.0 \times 10^{-3}$ | 15 | 4.8 | $1.0 \times 10^{-3}$ |
| Inds | 76 | 0.15 | $1.9 \times 10^{-3}$ | 0.86 | 33 | 25 | $1.0 \times 10^{-3}$ |
| Side:Inds | 76 | $1.4 \times 10^{-2}$ | $1.9 \times 10^{-4}$ | $8.4 \times 10^{-2}$ | 3.2 | 28 | $1.0 \times 10^{-3}$ |
| Residuals | 154 | $9.1 \times 10^{-3}$ | $5.9 \times 10^{-5}$ | $5.3 \times 10^{-2}$ | | | |
| Total | 307 | 0.17 |  |  |  |  |  |

